# The evolutionary dynamics of locally adaptive chromosome inversions in *Mimulus guttatus*

**DOI:** 10.1101/2023.12.06.570460

**Authors:** Leslie M. Kollar, Lauren E. Stanley, Sunil K. Kenchanmane Raju, David B. Lowry, Chad E. Niederhuth

## Abstract

Chromosomal inversion polymorphisms are ubiquitous across the diversity of diploid organisms and play a significant role in the evolution of adaptations in those species. Inversions are thought to operate as supergenes by trapping adaptive alleles at multiple linked loci through the suppression of recombination. While there is now considerable support for the supergene mechanism of inversion evolution, the extent to which inversions trap pre-existing adaptive genetic variation versus accumulate new adaptive variants over time remains unclear. In this study, we report new insights into the evolutionary dynamics of a locally adaptive chromosomal inversion polymorphism (inv_chr8A), which contributes to the evolutionary divergence between coastal perennial and inland annual ecotypes of the yellow monkeyflower, *Mimulus guttatus*. This research was enabled by the sequencing, assembly, and annotation of new annual and perennial genomes of *M. guttatus* using Oxford Nanopore long-read sequencing technology. In addition to the adaptive inv_chr8A inversion, we identified three other large inversion polymorphisms, including a previously unknown large inversion (inv_chr8B) nested within the inv_chr8A. Through population genomic analyses and comparative genomics, we determined that the nested inv_chr8B inversion is significantly older than the larger chromosomal inversion in which it resides. We also evaluated key candidate genes involved in gibberellin biosynthesis and anthocyanin regulation, which we hypothesize to underlie the adaptive phenotypic effects of the inv_chr8A inversion. Although little evidence was found to suggest that inversion breakpoint mutations drive adaptive phenotypic effects, our findings support the supergene mechanism of adaptive evolution as a dynamic and continuous process.

## Introduction

Chromosomal inversions have been implicated in evolutionary adaptations since classic studies in *Drosophila* found that inversion polymorphisms are frequently correlated with environmental conditions (Adrion et al., 2015; Dobzhansky, 1951, 1970; Hoffmann & Rieseberg, 2008; Kapun & Flatt, 2019; Kirkpatrick & Kern, 2012; Villoutreix et al., 2021). Large chromosomal inversions rearrange the structure of genomes causing major phenotypic consequences (Dobzhansky, 1936; Elgin & Reuter, 2013; Lupski, 1998; Marshall et al., 2008; Muller, 1930; Puig et al., 2015). Inversions strongly suppress genetic recombination in heterokaryotic individuals (inversion heterozygotes) because recombinant gametes are unbalanced (Dobzhansky, 1970; Huang & Rieseberg, 2020; Rieseberg, 2001). Researchers have thus hypothesized that inversions could act as supergenes by holding together haplotype blocks containing multiple locally adaptive polymorphisms through suppressed recombination (Berdan et al., 2023; Darlington & Mather, 1950; Dobzhansky, 1970; Kirkpatrick & Barrett, 2016; Kirkpatrick & Barton, 2006; Kirkpatrick & Kern, 2012; Schwander et al., 2014; Thompson & Jiggins, 2014).

The realization that inversions could spread rapidly as a result of trapping locally adaptive alleles at linked loci led Kirkpatrick and Barton to state a decade ago: “If the local adaptation mechanism is so powerful, why are inversions not everywhere? One possibility is that they in fact are, and that their frequency has been greatly underestimated” (Kirkpatrick & Barton, 2006). Since Kirkpatrick and Barton’s 2006 paper, it has become clear that inversion polymorphisms truly are everywhere and are frequently associated with evolutionary adaptations. At least 49 unique inversions have been linked to adaptive phenotypic traits in *Anopheles* mosquitoes (D. Ayala et al., 2014). Inversions have been shown to control the phenotypes underlying mimicry in butterflies (Joron et al., 2011; Kunte et al., 2014; Thompson & Jiggins, 2014) as well as mating and migratory behaviors in birds (Jeong et al., 2022; Lamichhaney et al., 2016; Lundberg et al., 2023; Tuttle et al., 2016). Shifts in the frequency of inversion polymorphisms have been linked to adaptation to forest versus prairie habitats in deer mice (Hager et al., 2022; Harringmeyer & Hoekstra, 2022) and to global climate change in *Drosophila* (Anderson et al., 2005; Rane et al., 2015). Inversions appear to play a prominent role in adaptation and domestication of crops, including maize, wheat, and peaches (Fang et al., 2012; Pyhäjärvi et al., 2013; Zhang et al., 2015; Zhou et al., 2021). Human and chimpanzee genomes differ by nine major chromosomal inversions and >100 smaller ones (Marquès-Bonet et al., 2004; Newman et al., 2005), which may suggest they played a key role in speciation from our closest relatives (F. J. Ayala & Coluzzi, 2005; Navarro & Barton, 2003). Despite the extensive progress in understanding adaptive inversion evolution, there is still much to learn about how inversions evolve and what genes contribute to their overall phenotypic effects (Berdan et al., 2023).

In this study, we evaluated the evolutionary dynamics of multiple chromosomal inversion polymorphism in the yellow monkeyflower, *Mimulus guttatus* (syn. *Erythranthe guttata*). This work builds on the previous discovery of a chromosomal inversion polymorphism on chromosome 8 underlying a quantitative trait locus (*DIV1*) in this system. This inversion has shown to be definitively linked to local adaptation through a field reciprocal transplant experiment (Lowry & Willis, 2010). While circumstantial evidence has been presented for the role of inversions in local adaptation in many other systems (Beardmore et al., 1960; Butlin & Day, 1984; Chouteau et al., 2017; Dobzhansky, T, 1947; Hager et al., 2022; Mérot et al., 2020), this is the only inversion polymorphism that has been definitively linked to local adaptation in the wild (Berdan et al., 2023). The inversion at the *DIV1* locus, which we henceforth refer to as inv_chr8A, has large phenotypic effects on a suite of adaptive traits associated with the transition from an annual to perennial life history, including flowering time, branching, allocation to vegetative growth, and herbivore resistance (Lowry et al., 2019; Lowry & Willis, 2010). The inv_chr8A inversion polymorphism is widespread across the geographic range of *Mimulus guttatus*, with one orientation of the inversion commonly found in perennial populations, and the other in annual populations (Lowry & Willis, 2010). The inversion most strikingly contributes to the divergence of coastal perennial populations, which experience cooler conditions and moisture supplied by summer sea fog, and nearby inland annual populations, which complete their life cycle quickly due to diminishing water availability during summer drought (Hall et al., 2010; Lowry et al., 2008).

Recently, Coughlin and Willis (2019) provided evidence to support the hypothesis that the inv_chr8A trapped pre-existing adaptive alleles in crosses between annual *M. guttatus* and a perennial sister species, *M. tilingii*, which share the same ancestral chromosomal inversion orientation. The orientation of inv_chr8A found in perennial *M. guttatus* is derived, even though perenniality itself is thought to be the ancestral state in this system (Coughlan & Willis, 2019; Friedman, 2014). Further, Coughlin and Willis (2019) were able to exploit free recombination within the inv_chr8A region to show that there are at least two quantitative trait loci (QTL) for morphological divergence in crosses between annual *M. guttatus* and perennial *M. tilingii*. While this result provides some evidence for the supergene hypothesis, the dynamics of how the inversion has evolved over time remain and the genes underlying the supergene effects of the inversion are still unknown. Identification of the candidate genes that cause the phenotypic effects of inversion polymorphisms in *M. guttatus* have been difficult because available genome assemblies derived from short read sequencing are fragmented (Hellsten et al., 2013) and were available only for ecotypes with the annual orientation. As a result, the full complement of genomic DNA and the inversion breakpoints of the adaptive inv_chr8A and other known inversions have remained unidentified.

Here, we report a set of important new discoveries regarding chromosomal inversion polymorphisms in *M. guttatus*, some of which play crucial roles in the local adaptation and differentiation of annual and perennial ecotypes within this species. This progress was made possible by the sequencing, assembly, and annotation of two new genomes for coastal perennial and inland annual lines of *M. guttatus* using Oxford Nanopore technologies that we report here. With these genomes, we were able to overcome previous challenges and identify the breakpoints of these inversions as well as the full complement of genes within the inversions, including inv_chr8A. To identify candidate genes for the adaptive phenotypic effects of the inversions, as well as explore possible phenotypic consequences of the inversion breakpoint mutations themselves, we combined our new genome assemblies with a set of existing population genetic and gene expression datasets. We also combined our analyses with a population genomic data set to evaluate whether particular genes and loci within the adaptive inv_chr8A inversion had undergone recent selective sweeps in either coastal or inland habitats. Through this process, we discovered another large chromosomal inversion (inv_chr8B) nested within inv_chr8A. This finding suggests that this supergene may have begun as a moderate sized inversion that then expanded by being trapped within a larger inversion. Overall, this study brings us closer to understanding how chromosomal inversions evolve and why genomes rearrange dramatically over long periods of evolutionary history.

## Materials and Methods

### Tissue harvesting and DNA extraction

Our genome sequencing efforts focused on one inland annual line, LMC-L1 (Yorktown, CA, 38° 51’ 50.3388’’ N, 123° 5’ 2.1012’’ W), and one coastal perennial line, SWB-S1 (Irish Beach, CA, 39° 2’ 9.5388’’ N, 123° 41’ 25.6812’’ W). While these lines are significant because they have been used in multiple studies (Friedman, 2014; Gould et al., 2018; Lowry et al., 2019; Lowry & Willis, 2010), we recognize that these lines are not representative of all coastal perennial and inland annuals. LMC-L1 was inbred for 6 generations and SWB-S1 was inbred for 14 generations in the Duke University greenhouses and the Michigan State University growth facilities. The fewer number of generations inbreeding of LMC-L1 is due to the line having a high level of sterility-based inbreeding depression. Floral buds were collected for DNA extractions from plants that were grown in growth chambers in the following conditions: 16 hour day length, 22°C day and 18°C night temperatures, with 60% relative humidity, and a light intensity of 460mE. DNA extracted from this tissue was used for both long read Nanopore and short read Illumina sequencing. For Nanopore sequencing, nuclei were extracted from the floral bud tissue following Lu et al (Lu et al., 2017), with minor modifications. We ground 1-1.5 g of floral bud tissue into a coarse powder in liquid nitrogen. Ground tissue was resuspended in LB01 buffer (15 mM Tris pH 7.5, 2 mM EDTA (Ethylenediaminetetraacetic acid), 80 mM KCl, 20 mM NaCl, 15 mM 2-Mercaptoethanol, 0.15% Triton-X 100 and 0.5 mM Spermine) on ice with intermittent mixing. Tissue homogenate was then filtered through four-layers of miracloth (EMD Millipore Corp) followed by filtration through a 20 μm cell strainer (pluriStrainer®). Filtrate was carefully pipetted on top of equal volume Density Gradient Centrifugation Buffer (1.7 M Sucrose, 10 mM Tris-HCl pH 8.0. 2 mM MgCl2, 5 mM 2-Mercaptoethanol, 1 mM EDTA, 0.15% Triton-X 100) in a 50 ml falcon tube and centrifuged at 2500g for 30 mins at 4C. After centrifugation, the supernatant was decanted and the nuclei pellet was used to isolate high-molecular weight (HMW) DNA using the Nanobind Plant Nuclei Big DNA kit (Circulomics, Baltimore, MD, Cat # NB-900-801-01).

### Oxford Nanopore PromethION sequencing

Large chromosomal rearrangements can be identified using population genetics methods with short-read sequencing (Todesco et al., 2020), but detecting the full complement of DNA within inversions and pinpointing inversion breakpoints requires long-read sequencing. To achieve this, we performed Oxford Nanopore Technologies (ONT) sequencing. Nanopore libraries were prepared using the Oxford Nanopore SQK-LSK110 Ligation Sequencing Kit and loaded onto a FLO-PRO002 (vR9.4.1) flow cell. Sequencing was carried out on a PromethION sequencer (21.02.7) for 96 hours, followed by basecalling with the Nanopore Guppy basecaller software (v4.3.4). We obtained 12,580,640 (120.74 GB) and 13,106,600 (133.36 GB) nanopore reads for LMC-L1 and SWB-S1, respectively, with median read lengths of 6.1 kb and 7.7 kb. Quality trimming resulted in 9,255,001 reads for LMC-L1 (337x coverage) and 9,423,361 reads for SWB-S1 (360x coverage), with 73.6% and 71.9% of reads passing filtering for LMC-L1 and SWB-S1, respectively.

### Illumina Sequencing

We polished the genome assembly and performed SNP calling using short-read sequencing. DNA was extracted from unopened floral buds using a DNeasy Plant Mini Kit (Qiagen) and prepped with the Illumina TruSeq Nano DNA Library prep kit. Sequencing was conducted on one lane of an Illumina NovaSeq 6000 S4 flow (2×150) using a NovaSeq V1.5 300-cycle reagent kit. Base calling was done with Illumina RTA v3.4.4, and the output was demultiplexed and converted to FastQ format using Bcl2fastq v2.20.0.

### Oxford Nanopore PromethION assembly, polishing, and scaffolding

Adapters were removed from the reads using Porechop v0.2.5 (https://github.com/rrwick/Porechop) and quality filtered using NanoLyse and Nanofilt (-q 0 and -l 300). We removed possible contaminants using a microbe genebank (genbank-k31.sbt.json). *De novo* genome assembly was conducted with Flye v2.9.1 (Kolmogorov et al., 2020), followed by two rounds of polishing using Racon (v1.5.0) (Vaser et al., 2017) and a single round of Medaka (https://github.com/nanoporetech/medaka). The genome was finished with two rounds of Pilon (Walker et al., 2014) polishing using the Illumina sequencing. We checked for misassembly using Tigment-long (Jackman et al., 2018). Scaffolding into chromosomes was initially performed using Allmaps (Tang et al., 2015) by combining genetic map data (Lowry & Willis, 2010) and synteny to the IM62, TOL, and NONTOL genomes (phytozome-next.jgi.doe.gov). Additional contigs were added to the chromosomes through an iterative process using ragtag (Alonge et al., 2022) and lrscaf (Qin et al., 2019), combined with manual inspection and correction. Genome assembly statistics were calculated using assembly-stats (*v1.0.1*, https://github.com/sanger-pathogens/assembly-stats). Completeness of the genome assembly was verified throughout the pipeline using BUSCO v2.0 (Simão et al., 2015) and the LTR Assembly Index (LAI) pipeline (Ou et al., 2018).

### Gene annotations

Genes were annotated using the MAKER pipeline (Cantarel et al., 2008; Holt & Yandell, 2011). Support for gene models was provided by transcripts assembled from RNA-seq and protein alignments. Illumina reads were trimmed for adapter sequences using trimmomatic (v0.39; (Bolger et al., 2014) and aligned to their respective genome using HISAT2 (v2.2.1; (Kim et al., 2019)). PacBio reads were aligned by minimap2 (v2.24; (Li, 2018; *Phytozome v13*, n.d.)). Transcripts were assembled by StringTie2 (v2.2.1; (Kovaka et al., 2019)). For protein alignments, protein sequences from *Arabidopsis thaliana* (Araport11;(Cheng et al., 2017), *Oryza sativa* (v7; (Kawahara et al., 2013)), *Solanum lycopersicum* (ITAG4.0; (Hosmani et al., 2019)), and UniProtKB/Swiss-Prot plants (release-2022_02; (Schneider et al., 2005)) were aligned to each genome using Exonerate protein2genome (Slater & Birney, 2005). Repeatmasker (v4.1.1; (*RepeatMasker Home Page*, n.d.)) was used to soft-mask the genome using TE sequences identified by Extensive *de-novo* TE Annotator (EDTA, see TE Annotations). An initial round of MAKER was run using the soft-masked genome and gff files from the transcriptome assembly and protein alignment. Annotations from this initial round of MAKER were then used to train SNAP (v2013_11_29; (Korf, 2004) and AUGUSTUS (v3.4.0; (Stanke & Morgenstern, 2005)). To prevent overfitting, only 600 randomly sampled MAKER annotations were used to train AUGUSTUS. MAKER was then run a second time using models from SNAP and AUGUSTUS. To identify putative genes missed by MAKER in either genome, we next used Liftoff (v1.6.3; (Shumate & Salzberg, 2020)) to transfer annotations from one genome onto the other. Lifted annotations were then filtered using GffCompare (v0.11.2; (Pertea & Pertea, 2020)) to remove annotations with significant overlap to existing MAKER annotations. Remaining lifted annotations with valid open reading frames (ORF; “valid_ORFs=1” or “valid_ORF=True” in the GFF attributes column) were considered as putative genes, while those without a valid ORF (“valid_ORFs=0” or “valid_ORF=False” in the GFF attributes column) were considered putative pseudogenes. This process was repeated using annotations from the reference IM62 (v2.0; Hellsten et al., 2013) genome annotation. Genes were flagged as potentially misannotated transposons as described previously (Bowman et al., 2017), with the exclusion of searching of Gypsy HMM profiles, as we found that this led to too many real genes being excluded. Given the often blurry line between genes and transposons, we retained these gene models in the annotation and simply flag them as possible transposons.

### Gene Function

Gene functions were assigned by first searching for PFAM domains using InterProScan (v5.57-90.0) and then identifying the top five hits with DIAMOND BLASTX against Arabidopsis TAIR10 proteins (e-value cutoff 1e-6). Results were integrated using a custom script from Kevin Childs (available at https://github.com/niederhuth/mimulus-assembly).

### Structural variant and SNP calling and annotation

Structural variants were called via the whole genome alignment approach using MUM&Co (*v3.0.0*, O’Donnell & Fischer, 2020). Pseudo-chromosome assemblies were aligned to each other, switching the reference and query, for calling SVs for each line. SVs (inversions, deletions, insertions, and duplications) were quality filtered to ≥ 50 bp. Because both SWB-S1 and LMC-L1 were inbred lines, we removed heterozygous variants as those are likely false positives. Lastly, we removed called insertions and deletions with a continuous strand of “NNN,” as these segments represent gaps created by RagTag and are likely false positives. Structural variants were annotated using BEDtools *intersect* (Quinlan & Hall, 2010) to identify overlapping genes and structural variants.

To call SNPs, we aligned the Illumina sequences to each genome. We quality trimmed using Trimmomatic v0.39 (Bolger et al., 2014) to a minimum length of 50 bps and a quality of phred33. We used BWA-MEM2 (Vasimuddin et al., 2019) to align SWB-S1 reads to the inland annual genome (98.63% alignment rate) and LMC-L1 reads to the coastal perennial (98.3% alignment rate) genome. SNPs were called using GATK (McKenna et al., 2010). We marked duplicates using PICARDTOOLS’ *MarkDuplicates* function and then used GATK’s *HaplotypeCaller* to call SNPs in individual samples. We then genotyped the SNPs using GATK’s *GenotypeGVCF* and filtered them using *VariantFiltration*.

SNPs were annotated using ANNOVAR (Wang et al., 2010) and filtered to include: frameshift insertions, frameshift deletions, stop loss, stop gain, and splicing. ANNOVAR (Wang et al., 2010) was used to identify synonymous and nonsynonymous SNPs. To estimate a rate of synonymous substitutions at 4-fold degenerate sites, we identified 4-fold degenerate sites using degenotate (https://github.com/harvardinformatics/degenotate.git). We intersected the synonymous SNPs with the list of 4-fold degenerate sites using bedtools *intersect (Quinlan & Hall, 2010)*.

We also reanalyzed SNPs from pooled sequencing from Gould et al (Gould et al. 2017) using our genome assemblies (SI Appendix, supplementary methods 2).

Copy number (CNVs) and presence absence variants (PAVs) were identified with a custom script using the outputs from GENESPACE. We included the *M. guttatus* IM62 v2 genome (Hellsten et al., 2013) as well as SWB-S1 and LMC-L1 genomes.

### Synteny, PAVs, CNVs, and inversion breakpoints

Our new genome assemblies fully localized breakpoint regions in the inv_chr8A inversion on chromosome 8. Due to the complex repetitive nature of the breakpoints, we defined them as regions extending from the last gene of the upstream collinear syntenic block to the second gene of the inverted block. Similarly, the breakpoint on the opposite end spans from the second-to-last gene of the inverted block to the first gene of the downstream collinear block. (SI Appendix, Fig. S13). To identify a stringent set of syntenic genes between ecotypes, we implemented GENESPACE (Lovell et al., 2022), a pipeline that uses both OrthoFinder (Emms, n.d.; Emms & Kelly, 2019) and blastp. For all estimates of sequence level differences, we used the first base pair of the first gene of the inverted syntenic block to the last gene of the inverted syntenic block. In addition to inv_chr8A, we were able to localize the boundaries of another previously known inversion on chromosome 5 (inv_chr5A), as well as two previously unknown large inversions and many small inversions. These newly identified inversions include a smaller inversion nested within inv_chr8A (inv_chr8B) and a large inversion on chromosome 14 (inv_chr14A). For all of these inversions, the repetitive nature of the region around the inversion breakpoints made it challenging to define the exact nucleotide of the breakpoint mutation. While we could not identify exact breakpoints, we could define which genes were closest to both sides of each breakpoint. Expression differences at the breakpoints were identified by reanalyzing Gould et al (Gould et al. 2018, SI appendix, supplementary methods 3).

### qRT-PCR of *GA20ox2* (AT5G51810, Migut.H00683)

To determine whether expression differences in *GA20ox2* might contribute to phenotypic differences between ecotypes, we conducted qRT-PCR because this gene is expressed at a level too low to be analyzed with RNA-seq. We extracted RNA from three biological replicates of leaf tissue and the floral shoot apex from wild-type LMC-L1 and SWB-S1 plants using the Spectrum Plant Total RNA Kit (Sigma). cDNA was synthesized for each sample using GoScript Reverse Transcription Mix, Oligo(dT) (Promega). We performed qRT-PCR using Power SYBR Green PCR master mix (Applied Biosystems) on a CFX96 touch real-time PCR machine (Bio-Rad). The PCR program was as follows: 40 cycles at 95°C for 15 s and 60°C for 30 s. We determined amplification efficiencies for each primer pair using a dilution series (1:2, 1:4, 1:8, 1:16) of pooled cDNA samples. We used *UBC*, the ubiquitin-conjugating enzyme gene, as a reference gene to calculate the relative expression of *GA20ox2* using the formula (E_ref_)^CP(ref)^/(E_target_)^CP(target)^ (Pfaffl, 2001). No differential expression was found for leaf tissue and these results are not reported (two-tailed Student’s t-test, p > 0.05).

## Results

### De novo assembly of M. guttatus inland annual and coastal perennial lines

The final, polished genome assemblies were approximately 277 Mbps (335X coverage ONT and 150X coverage WGS) spanning 1,198 contigs for LMC-L1 (inland annual) and 278 Mbps (365X coverage ONT and 151X) spanning 707 contigs for SWB-S1 (coastal perennial). Both assemblies demonstrated high contiguity with a contig N50 of 5.83 Mbps (LMC-L1) and 4.90 Mbps (SWB-S1) and a scaffold N50 of 18.2 Mbps (LMC-L1) and 18.8 Mbps (SWB-S1). Based on genetic maps and synteny, we assembled 93.0% of the LMC-L1 bps and 93.7% of the SWB-S1 sequence into 14 chromosomes (Hellsten et al., 2013). Our genomes recovered more than 98% of the eudicot set of BUSCO (Simão et al., 2015) orthologs (eudicots_odb10), an improvement from 96.9% in the previously available IM62 v2 assembly (Hellsten et al., 2013). The LTR Assembly Index (LAI), an independent assessment of assembly quality based on the completeness of assembled LTR retrotransposons (Ou et al., 2018), was much higher in both LMC-L1 (13.47) and SWB-S1 (16.28) than the IM62 v2 reference (8.79), likely indicating more complete assembly of repetitive regions (S1 Appendix, Table S1).

We annotated a total of 27,583 genes in LMC-L1 and 26,876 genes in SWB, in comparison to 28,140 genes in IM62 v2 (Hellsten et al., 2013) (SI Appendix, Table S1). We compared the LMC-L1 and SWB-S1 genomes using *GENESPACE* (Lovell et al., 2022), identifying 19,245 syntenic genes between LMC-L1 and SWB-S1 (SI Appendix, Table S2). Additionally in LMC-L1, we found 5,105 presence absence variants (PAVs), 3,026 pseudogenes, and 1,411 tandem duplicates. In SWB-S1, we found 4,707 PAVs, 3,110 pseudogenes, and 1,401 tandem duplicates.

### Genome wide SNPs and structural variants

To identify larger structural variants (SVs) between the LMC-L1 and SWB-S1 genomes, we used *MUM&CO,* a whole genome alignment method (O’Donnell & Fischer, 2020). The number of SVs between the LMC-L1 and SWB-S1 genomes are reported in SI Appendix, Table S3. Across the genome there were 12,013 genes associated with structural variants in LMC-L1 (variants affecting genes: 1,379 insertions, 8,493 deletions, 72 duplications, and 2,069 inversions) and 11,582 genes associated with structural variants in the SWB-S1 (variants affecting genes: 1,375 insertions, 8,300 deletions, 51 duplications, and 1,856 inversions).

To identify SNPs between the LMC-L1 and SWB-S1 accessions, we aligned whole genome sequencing (WGS) from one accession to the other accession’s reference genome. From these alignments, we focused on the evolution of coding regions by identifying high impact SNPs, which include the following variant types: stop gain (2,787 in SWB-S1 and 2,638 in LMC-L1), stop loss (723 in SWB-S1 and 700 in LMC-L1), frameshift deletion (7,299 in SWB-S1 and 7,541 in LMC-L1), frameshift insertion (7,002 in SWB-S1 and 6,951 in LMC-L1), and splicing (3,986 in SWB-S1 and 3,392 in LMC-L1). We also identified all other synonymous and nonsynonymous SNPs. All coding region SNPs are reported for both genomes in SI Appendix, Tables S4 and S5.

### Identification of large chromosome inversions and their breakpoints

The assembly of the nanopore long-read genomes allowed us for the first time to identify large chromosomal inversions from sequencing data alone as well as identify the locations of the breakpoints mutations that formed these inversions in the first place. The locally adaptive Inv_chr8A was found to span 6.7 Mb in LMC-L1 and 5.6 Mb in SWB-S1 (Fig. 1A, SI Appendix Fig. S1 and Table S5). In addition, our analysis led to the discovery of large inversion nested within inv_chr8A. This nested inversion(inv_chr8B) spans 213,792 bps in LMC-L1 and 261,302 bps in SWB-S1 (Fig. 1B, SI Appendix Fig. 1 and Table S5). Beyond chromosome 8, we were able to localize a previously identified inversion on chromosome 5 (inv_chr5A), which is approximately 4.2 Mb long in LMC-L1 and 4.0 Mb in SWB-S1 (Fig. 1A, SI Appendix Fig. 1 and Table S5). While we have successfully identified inv_chr5A from our genome assemblies, it is located near the end of the chromosome arm and there are gaps within the inversion in LMC-L1 of the assemblies. Therefore, there is uncertainty about the location of this inversion’s breakpoint toward the end of the chromosome. Finally, we identified a fourth large inversion on chromosome 14 (inv_chr14A), which was previously unknown. The inv_chr14A inversion has an approximate size of 2.4 Mb in LMC-L1 and 2.8 Mb in SWB-S1 (Fig. 1A, SI Appendix, Fig 1 Table S5).

**Figure 1.**
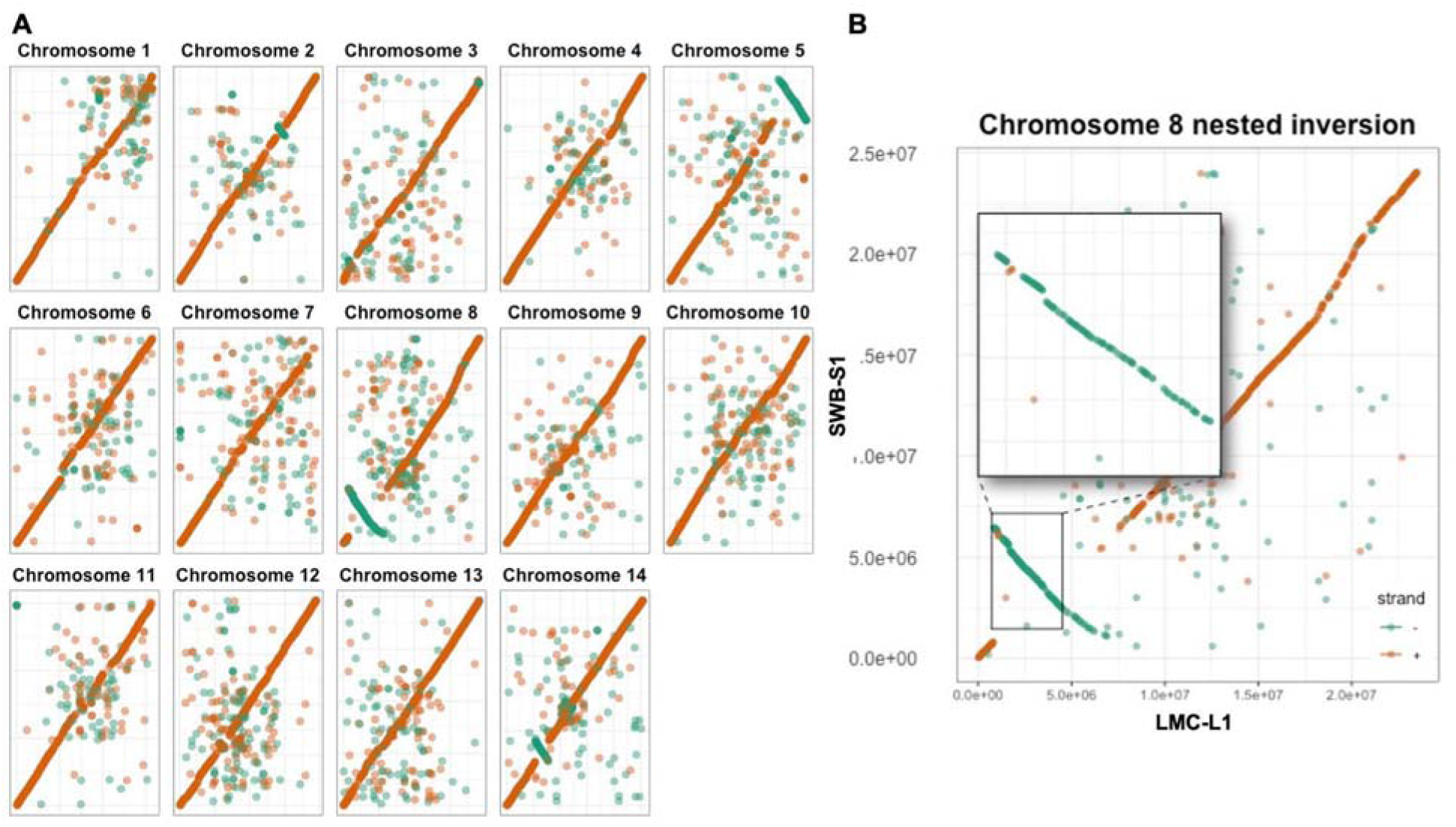
**A**. Whole genome alignment of all 14 chromosomes between the SWB-S1 and LMC-L1 genomes. Large inversions on chromosomes 5, 8, and 14 are visible in the dotplots, where the teal dots represent the negatively oriented strands while orange represent positively oriented strands. **B.** The inv_chr8B captured within inv_chr8A.

Because of the differences in size of the inversion regions in the two genome assemblies, we were curious whether there were any major differences in gene content between the inversion orientations. Our analyses revealed striking differences in gene content. Inv_chr8A (including inv_chr8B) had 177 LMC-L1-specific genes, 125 SWB-S1-specific genes, and 613 genes shared between the two genomes. Inv_chr8B shares 36 genes between the genomes but has 9 LMC-L1-specific genes and 8 SWB-S1-specific genes. Inv_chr5A had a total of 113 LMC-L1 specific genes and 90 SWB-S1 specific genes but shared 396 genes between the LMC-L1 and SWB-S1. Lastly, the chromosome 14 inversion had 247 LMC-L1-specific genes, 308 SWB-S1-specific genes, and 1,133 shared genes. Many of these large gene content differences are due to PAVs and copy number variants (CNVs). Errors in gene annotations could also contribute to PAVs and CNVs and thus, our estimated gene content differences.

While previous studies have demonstrated the role of inv_chr8A in local adaptation (Lowry & Willis, 2010), and implicated inv_chr5A (Gould et al., 2017), our improved assemblies now capture the full complement of genes within these inversions and make it possible to identify candidate genes that could have driven the geographic distributions of these inversions. In the following section, we highlight genetic differences between inversion orientations. CNVs, including PAVs, could explain both differences in inversion size and phenotypic differences. Within inv_chr8A, there were 338 CNVs with 302 being PAVs; in inv_chr8B, there were 19 CNVs with 17 being PAVs. Within inv_chr5A, there were 227 CNVs, with 203 being PAVs. In inv_chr14A, there were 100 CNVs, with 84 being PAVs. Some of these CNVs in the inversions could be explained by genes becoming pseudogenized within the inversion. In inv_chr8A, including inv_chr8B, there were 90 pseudogenes in LMC-L1 and 74 pseudogenes in SWB-S1 (SI Appendix, Table S6). Inv_chr8B included 6 pseudogenes in LMC-L1 and 8 pseudogenes in SWB-S1 (SI Appendix, Table S6). In inv_chr5A, there were 54 pseudogenes in LMC-L1 and 43 pseudogenes in SWB-S1 (SI Appendix, Table S6). Lastly, in inv_chr14A there were 22 pseudogenes in LMC-L1 and 19 pseudogenes in SWB-S1 (SI Appendix, Table S6). We report SNPs (all SNPs, high impact SNPs, and synonymous and nonsynonymous SNPs) falling within the chromosome inversions in SI Appendix, Tables S4 and S5. These SNPs were identified from WGS comparison of the SWB-S1 and LMC-L1 genomes.

### Evolutionary histories of large inversions

To determine which inversion orientations are ancestral, we implemented GENESPACE (Lovell et al., 2022) using *Mimulus lewisii,* a distant relative, as the reference genome for comparison (http://mimubase.org/FTP/Genomes/) (SI Appendix, Fig. S2). From this analysis, we reasoned that inversions sharing the same orientation as *M. lewisii* are in the ancestral orientation. We found the orientations of inv_ch5A (SI Appendix, Fig. S3), inv_chr8A (SI Appendix, Fig. S4), and inv_chr8B (SI Appendix, Fig. S4) are ancestral in LMC-L1. SWB-S1 has the ancestral orientation for inv_chr14A (SI Appendix, Fig. S5). The results for inv_chr8A are consistent with the results found by Coughlan et al. (Coughlan et al., 2021).

In addition to identifying which inversion orientation is ancestral, we were interested in the relative ages of inversion polymorphisms. Here, we compared the level of divergence between the LMC-L1 and SWB-S1 genomes for inverted regions and to the rest of the genome. When an inversion first forms, the two orientations of an inversion should have similar levels of divergence as the rest of the genome. As an inversion ages, each orientation should accumulate genetic changes that lead to elevated levels of divergence. To evaluate the relative ages of each inversion, we aligned Illumina reads from LMC-L1 to the SWB-S1 genome and aligned reads from SWB-S1 to the LMC-L1 genome. We then compared the levels of divergence at 4-fold degenerate synonymous sites for each inversion. The genome-wide average divergence at 4-fold degenerate sites was 0.043, regardless of which genome was used as the reference (SI Appendix, Table 2). Both the inv_chr5A (LMC-L1 Illumina resequencing: 0.055, SWB-S1 Illumina resequencing: 0.061) and inv_chr8A (LMC-L1 Illumina resequencing: 0.057, SWB-S1 Illumina resequencing: 0.058) inversions had elevated divergence relative to the genome-wide average, which suggests that these inversions have been segregating within this species for a long time. The inv_chr14A had similar levels of 4-fold degenerate site divergence (LMC-L1 Illumina resequencing: 0.041, SWB-S1 Illumina resequencing: 0.037) as the genome-wide average, suggesting that it is a relatively new inversion. Interestingly, the smaller nested inversion (inv_chr8B) had very high levels of divergence (LMC-L1 Illumina resequencing: 0.084 SWB-S1 Illumina resequencing: 0.083, SI Appendix, Table 2), which suggests that it is much older than any of the other inversions.

In addition to directly comparing divergence between the SWB-S1 and LMC-L1 genomes, we were able to evaluate whether there were differences in ecotype-wide polymorphism using a previous population genetic dataset (Gould et al., 2017). That study compared population genetic parameters for whole genome pooled sequences of 101 coastal perennial accessions versus 92 inland annual accessions (Gould et al., 2017). We reanalyzed this data using our new LMC-L1 and SWB-S1 genome assemblies (SI Appendix, Methods). Given the improvements in the genome assemblies over the previous reference IM62 v2 genome (Hellsten et al., 2013), we anticipated potentially new patterns in sequence diversity and differentiation. This new analysis also allowed us to evaluate how reference bias in alignments can affect population genetic summary statistic estimation. Regardless of the reference genome, the inland annual ecotype had higher nucleotide diversity (π = 0.040 regardless of reference genome). In contrast, the coastal ecotype pool differed slightly in diversity based on reference genome (π = 0.032 aligned to SWB-S1, π = 0.034 aligned to LMC-L1). The lower diversity of the coastal ecotype is consistent with prior studies and the hypothesis that the coastal ecotype was founded through a population bottleneck (Gould et al., 2017; Lowry et al., 2008) (SI Appendix, Fig. S6).

The population genetic data (Gould et al. 2017) allowed us to better evaluate the recent evolutionary history of the four large inversions. First, we compared the allelic differentiation (*G* statistic) for each inversion versus the rest of the genome. Consistent with our analysis of four-fold degenerate divergence between the SWB-S1 and LMC-L1 genomes, the nested inv_chr8B inversion had the highest level of allelic differentiation between the ecotypes (Fig. 2, SI Appendix, Fig. S7 and Table 1). The inv_chr8A and inv_chr5A inversions also had higher allelic divergence than the genome-wide average. Also consistent with the divergence result, the inv_chr14A inversion had similar levels of allelic differentiation as the rest of the genome. To evaluate whether any of the inversions had undergone a recent selective sweep, we compared the within ecotype diversity (π) for the inverted region to the rest of the genome. The idea here is that a significant reduction of within ecotype diversity for any of the inversions would be consistent with a recent selective sweep. To simplify this analysis, we calculated the ratio of π (π inland annual pool/π coastal perennial pool) for windows across the genome. A π-ratio elevated above the genome-wide average (π-ratio = 1.25) would be consistent with a sweep within the coastal perennial ecotype, while a reduction in the π-ratio would be consistent with a sweep in the inland annual ecotype. Overall, the distribution of π-ratios was similar for inv_chr5A, inv_chr8A, and inv_chr14A (Fig. 2, SI Appendix, Fig. S6 and Table 1). In contrast, the distribution of the π-ratio was significantly elevated for the nested inv_chr8B inversion, suggesting the possibility of a recent selective sweep in the coastal perennial ecotype.

**Figure 2.**
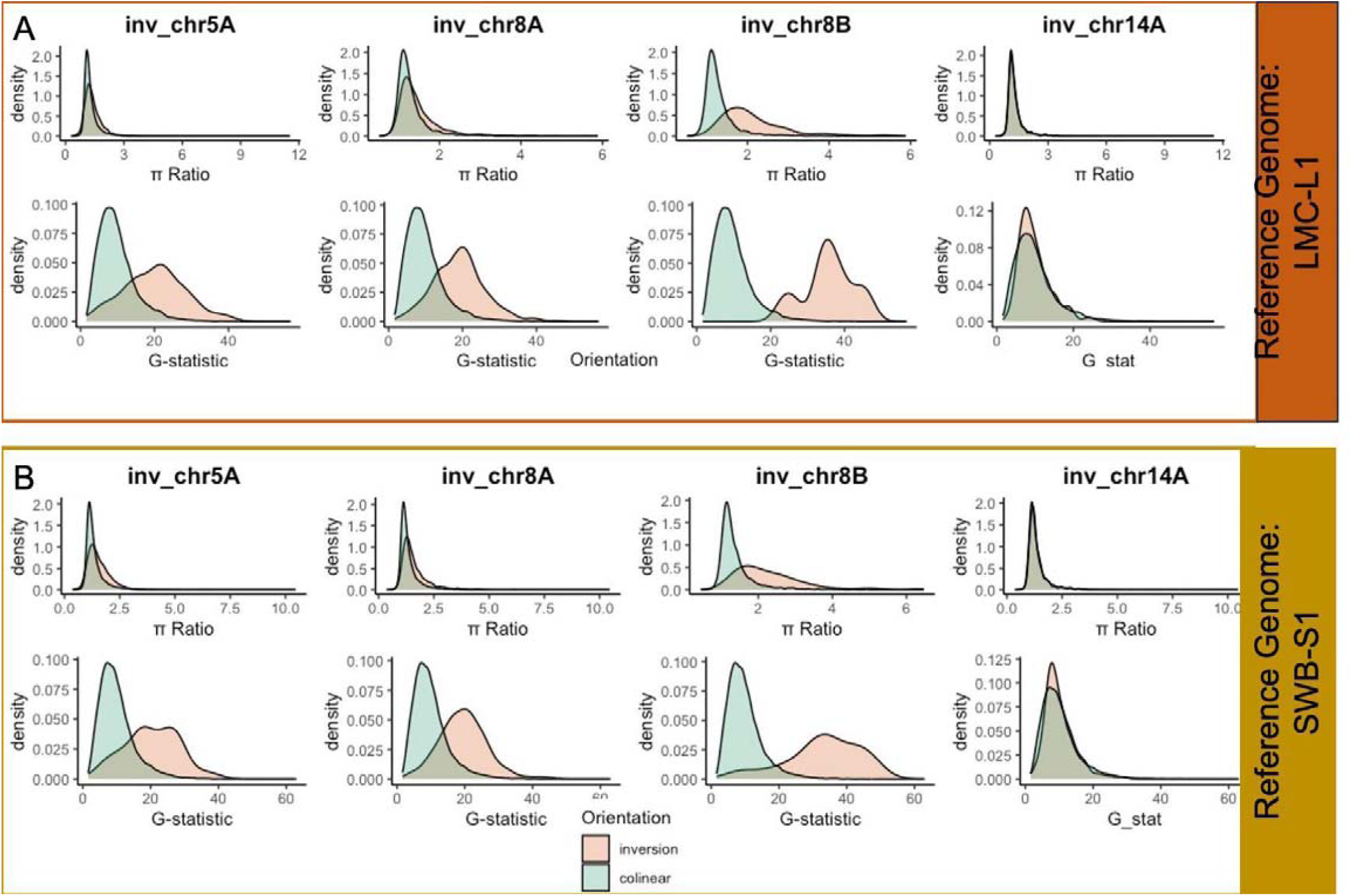
*G* statistic and π ratio for each of the inversions (salmon) compared to the non-inverted (teal) sequence of the same chromosome when aligned to the LMC-L1 Genome **A** and to the SWB-S1 genome **B**.

**Table 1.** Comparison of π and *G* statistic across the genome and for each inversion. IA represents the inland annual pool and CP represents the coastal perennial pool.

| Reference genome | Location | $\pi$ IA | $\pi$ CP | $\pi$ ratio | G statistic |
| --- | --- | --- | --- | --- | --- |
| SWB-S1 | Colinear regions of chromosome 5 | 0.040 | 0.032 | 1.26 | 9.98 |
| SWB-S1 | inv_chr5A | 0.040 | 0.029 | 1.41 | 20.28 |
| SWB-S1 | Colinear regions of chromosome 8 | 0.040 | 0.032 | 1.25 | 9.90 |
| SWB-S1 | inv_chr8A | 0.041 | 0.028 | 1.48 | 20.07 |
| SWB-S1 | inv_chr8A (without inv_chr8B) | 0.041 | 0.028 | 1.46 | 19.57 |
| SWB-S1 | inv_chr8B | 0.042 | 0.021 | 2.02 | 32.88 |
| SWB-S1 | Colinear regions of chromosome 14 | 0.040 | 0.032 | 1.26 | 10.16 |
| SWB-S1 | inv_chr14A | 0.041 | 0.033 | 1.25 | 10.09 |
| SWB-S1 | genome-wide | 0.040 | 0.032 | 1.26 | 10.16 |
| LMC-L1 | Colinear regions of chromosome 5 | 0.043 | 0.037 | 1.18 | 8.19 |
| LMC-L1 | inv_chr5A | 0.040 | 0.031 | 1.28 | 20.05 |
| LMC-L1 | Colinear regions of chromosome 8 | 0.040 | 0.033 | 1.21 | 10.70 |
| LMC-L1 | inv_chr8A | 0.040 | 0.029 | 1.37 | 19.64 |
| LMC-L1 | inv_chr8A (without inv_chr8B) | 0.040 | 0.030 | 1.35 | 18.71 |
| LMC-L1 | inv_chr8B | 0.042 | 0.021 | 1.93 | 35.76 |
| LMC-L1 | Colinear regions of chromosome 14 | 0.039 | 0.031 | 1.23 | 11.03 |
| LMC-L1 | inv_chr14A | 0.041 | 0.034 | 1.20 | 9.83 |
| LMC-L1 | genome-wide | 0.040 | 0.034 | 1.20 | 10.01 |

**Table 2.** LMC-L1 and SWB-S1 SNPs called from WGS were annotated and filtered to include synonymous SNPs at 4-fold degenerate sites for each inversion and genome-wide. Note SNPs for LMC-L1 were called after aligned WGS data to the SWB-S1 genome and SWB-S1 SNPs were called after aligned WGS data to the LMC-L1 genome.

| Genotype | Location | 4-fold degenerate SNPs | 4-fold degenerate sites | <u>4-fold degenerate SNPs</u><br>4-fold degenerate sites |
| --- | --- | --- | --- | --- |
| LMC-L1 | Genome-wide | 174,745 | 4,106,234 | 0.043 |
| LMC-L1 | inv_5A | 4,497 | 81,202 | 0.055 |
| LMC-L1 | inv_8A | 8,030 | 139,941 | 0.057 |
| LMC-L1 | inv_8B | 636 | 7,569 | 0.084 |
| LMC-L1 | inv_14A | 1,634 | 39,641 | 0.041 |
| SWB-S1 | Genome-wide | 176,470 | 4,093,796 | 0.043 |
| SWB-S1 | inv_5A | 4,772 | 78,850 | 0.061 |
| SWB-S1 | inv_8A | 7,954 | 136,834 | 0.058 |
| SWB-S1 | inv_8B | 630 | 7,633 | 0.083 |
| SWB-S1 | inv_14A | 1,555 | 42,095 | 0.037 |

### Candidate genes for inversion supergene effects

With clear resolution of the breakpoints for inv_chr8A and inv_chr8B, we were able to localize candidate genes that could underlie the phenotypic effects of inversions. We identified genes affected by divergence as those with outlier (top 1%) windows for the *G* statistic (explained above). Across the genome, the top windows for *G* contained 753 LMC-L1 and 635 SWB-S1 genes and/or their promoters and transcription start site (TSS) regions or 3’ untranslated regions (UTR). Many of these outlier genes, promoters and TSS regions, and 3’ UTRs were syntenic between the LMC-L1 and SWB-S1 genomes, including 193 genes, 57 promoters and TSS regions, and 90 3’ UTRs. Some of these outlier regions also fall within the chromosome inversions. Outlier windows falling within genes, promoters and TSS regions, and 3’ UTRs are interesting because these could contribute to phenotypic differences between the ecotypes. While these results do not suggest a direct cause in phenotypic variation, it does suggest these outlier genes are differentiated and warrant further investigation.

One such gene is *GA20ox2*, which codes for a structural gene in the gibberellin hormone synthesis pathway. Foliar application of gibberellin growth hormone (GA3) to coastal perennial plants has been shown to increase internode elongation and convert branches from vegetative stolons to aerial flowering branches in a manner that mimics inland annual growth architecture (Lowry et al. 2019). *GA20ox2* falls within inv_chr8A and may contribute to adaptive divergence between the ecotypes. *GA20ox2* (MgS1_08g02280) was an outlier for *G* and π ratio in our reanalysis using the SWB-S1 reference genome, similar to what was found in Gould et al. (Gould et al., 2017) when using IM62 as the reference genome. Both MgS1_08g02280 and MgL1_08g0964, the gene names for *GA20ox2* in each of the two reference genomes, appear to be intact genes with ORFs. This was confirmed by Sanger sequencing, and both genes are expressed based on qRT-PCR (Fig. 3). In LMC-L1, expression of *GA20ox2* was ∼1.7-fold higher in floral shoot apex compared to SWB-S1.

**Figure 3.**
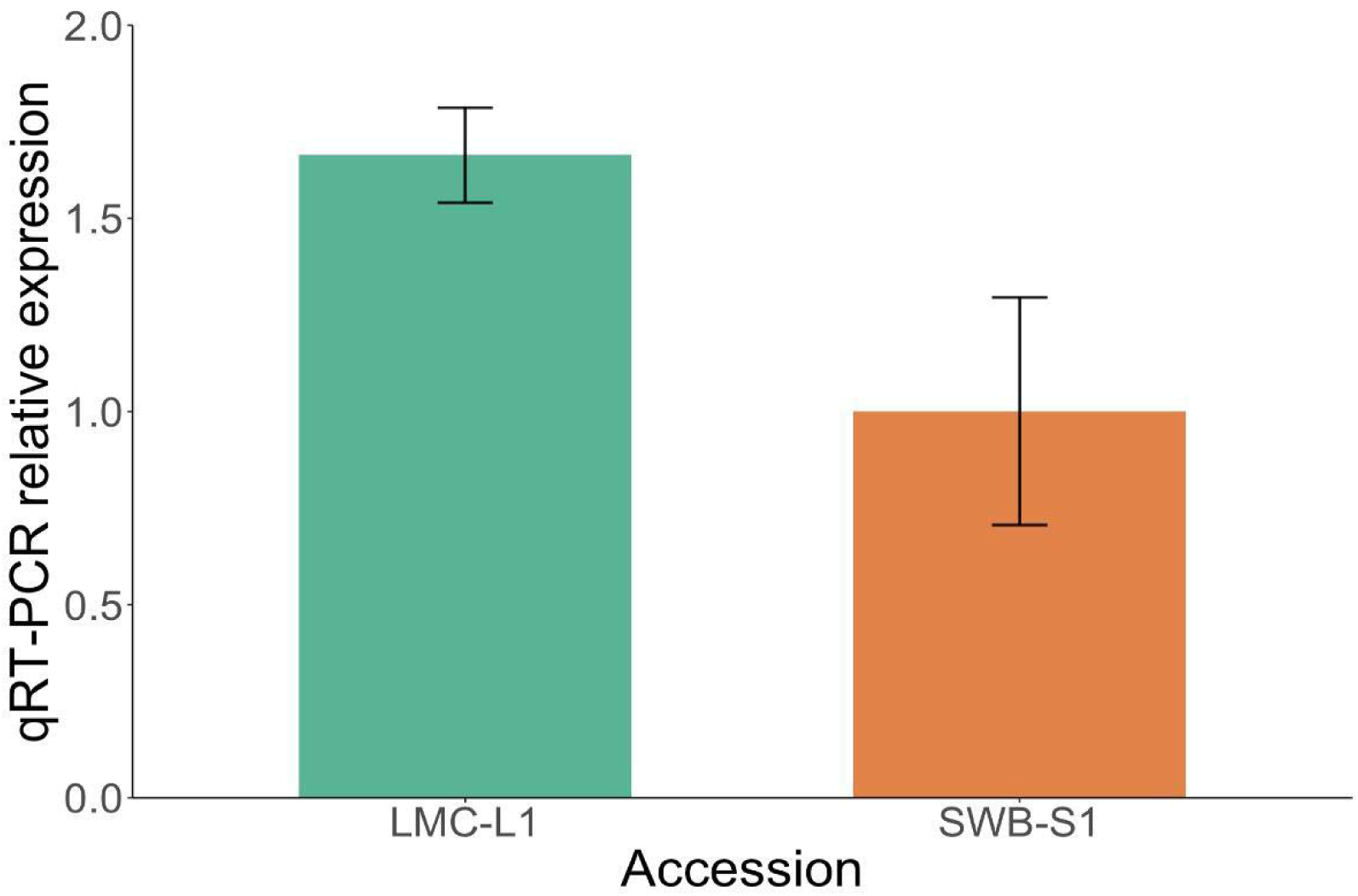
Quantitative RT-PCR showing expression of *GA20ox2* in wild-type LMC-L1 and SWB-S1 floral shoot apex (three biological replicates). Error bars show standard deviation.

Another likely set of genes underlying phenotypes that differentiate ecotypes are a tandem array of *R2R3-MYB* genes within inv_chr8A (Cooley et al., 2011; Lowry et al., 2012). Previous work showed that several calyx, corolla, and leaf anthocyanin pigmentation traits differentiating the ecotypes also map to the region containing inv_chr8A (Lowry et al., 2012). It has been difficult to resolve the number of tandem copies of *R2R3-MYB* with previously available genome assemblies (i.e. IM62, TOL, and NONTOL; phytozome-next.jgi.doe.gov/). Based on our CNV analysis, LMC-L1 and SWB-S1 have different numbers of copies of these subgroup six *R2R3-MYB* genes. The LMC-L1 genome has five partial or complete copies of this gene, with three being flagged as pseudogenes (genes without valid open reading frames) in our analysis, while the SWB-S1 genome appears to have six partial or complete copies of this gene, none of which were called as pseudogenes (Fig. 4). Interestingly, the subgroup six *R2R3-MYB* genes appear syntenic with conserved directionality in LMC-L1 and SWB-S1, except for MgL1_08g06980, which is in the opposite orientation of all the other MYBs. This could indicate that it was duplicated independently. In investigating why some LMC-L1 copies were called as pseudogenes, we found that MgL1_08g07004 matched perfectly to an annotation in IM62 (H00280), which is not annotated as an R2R3-MYB gene because it is missing distinguishing domains. However, MgL1_08g07004 almost perfectly matches MgL1_08g07000, especially in its intronic sequence, indicating that it is a partially duplicated gene.

**Figure 4.**
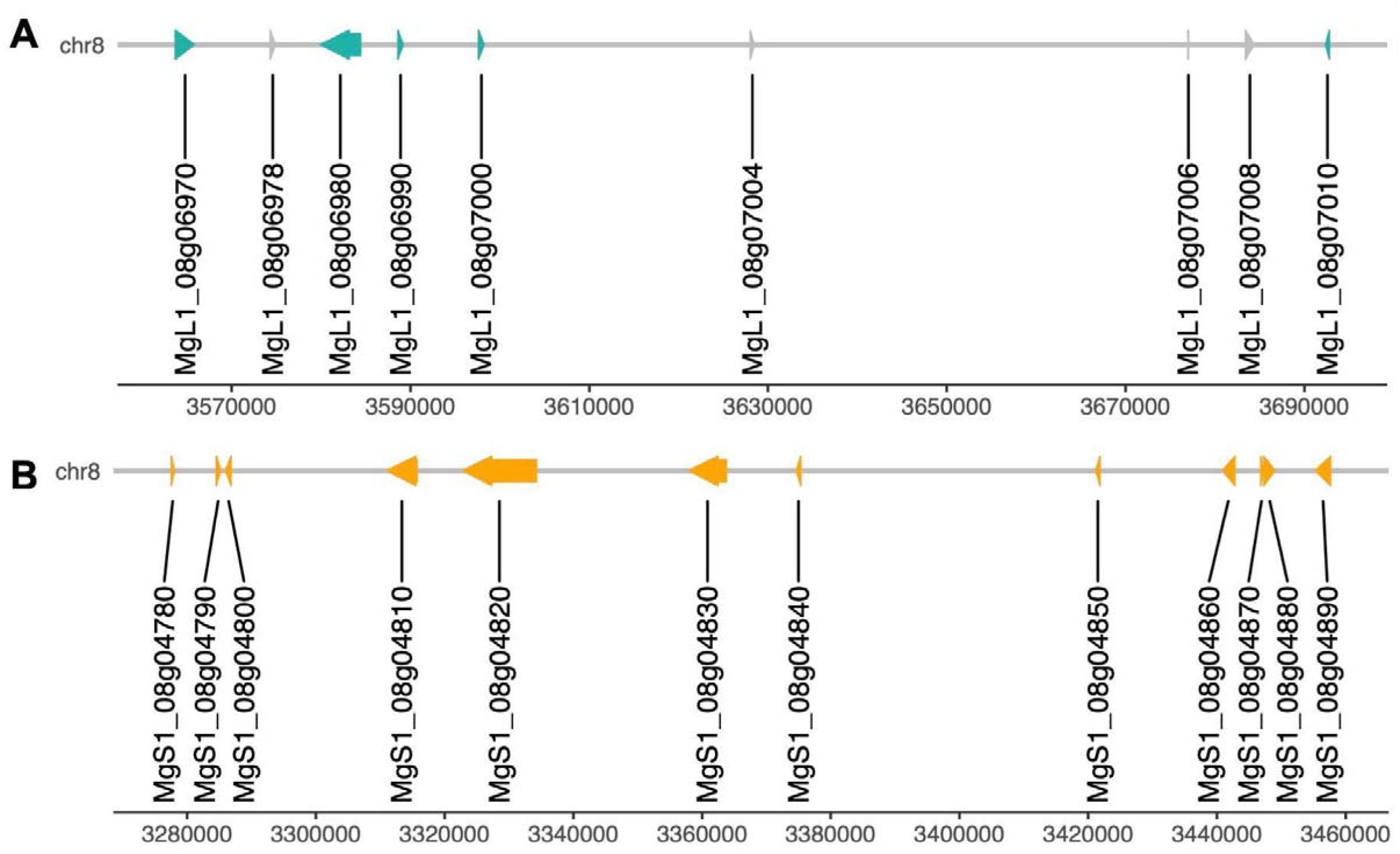
The MYB region along chromosome 8 in LMC-L1 (**A**) and SWB-S1 (**B**). Each arrow (teal in LMC-L1 and orange in SWB-S1) represents a gene and orientation. The gray arrows represent pseudogenes.

### Genes near inversion breakpoints were differentially expressed

The identification of inversion breakpoints allows for the evaluation of whether those breakpoints have disrupted genes in ways that could contribute to phenotypic effects. If an inversion breakpoint occurs within a gene, it would disrupt and likely eliminate its function. Breakpoints occurring near genes can also cause changes in gene expression by disrupting or changing the position of *cis*-regulatory elements. To establish whether gene expression might have been disrupted by the inversion breakpoint mutations, we evaluated transcript abundances within the breakpoint regions. We focused on the first two and last two genes of each inversion. We also evaluated transcript abundance for the first genes on either side just outside of the inverted region (SI Appendix, Fig. S13). We investigated these breakpoint region genes by looking at differentially expressed genes from RNAseq data (Gould et al., 2018) generated from a reciprocal transplant experiment (Fig. 5, SI Appendix, Table S7). We found that a total of 7 genes within the breakpoint regions across 4 inversions showed expression level differences.

**Figure 5.**
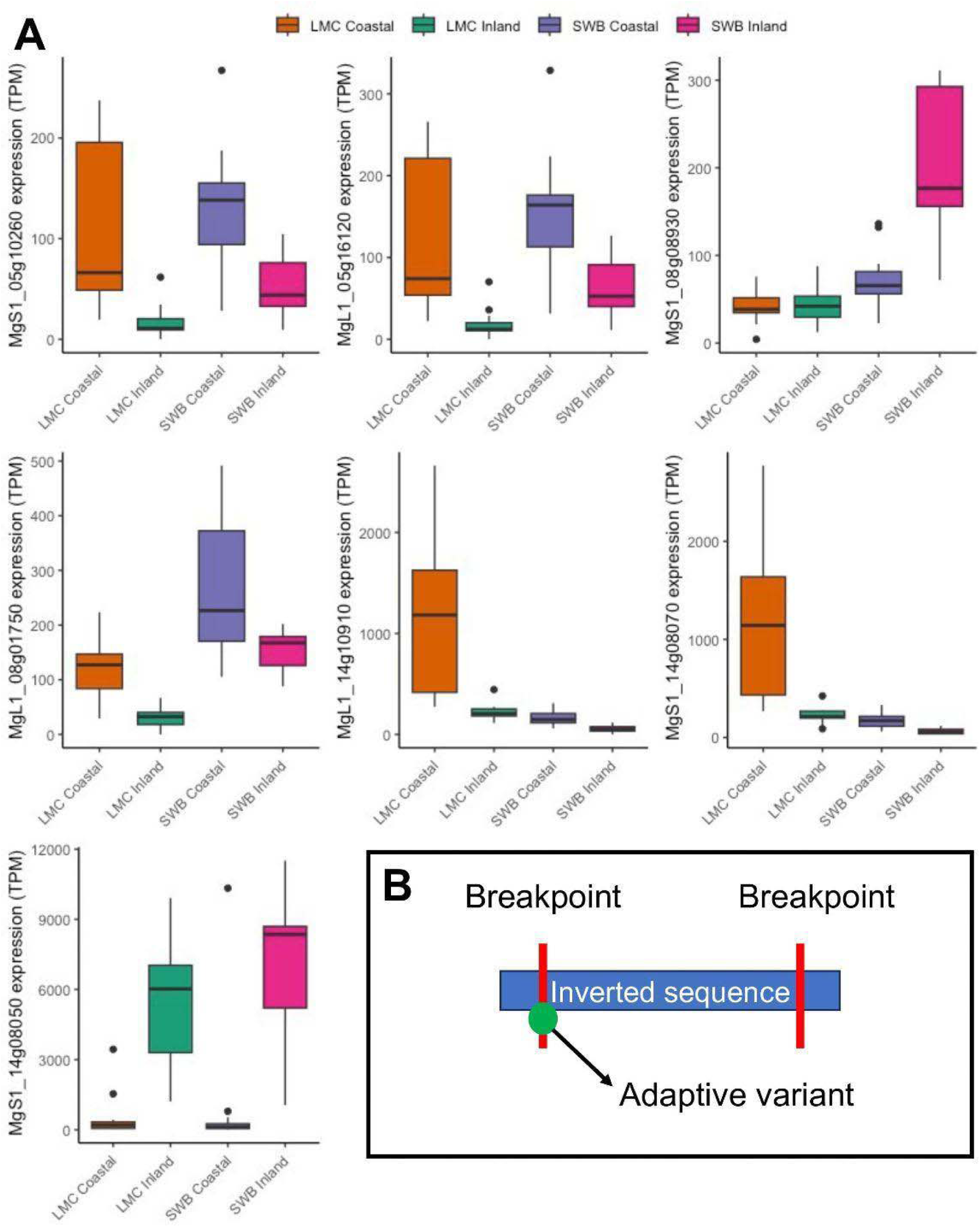
**A.** Differential expression represented in transcripts per million (TPM) in genes at the inversion breakpoints. Note, some plots are nearly identical because they are LMC-L1 (labeled LMC) and SWB-S1 (labeled SWB) orthologs (MgL14g10910 and MgS1_14g08070, MgS1_05g10260 and MgL1_05g16120). **B.** A diagram depicting the inversion breakpoint hypothesis.

## DISCUSSION

While chromosomal inversions are now well recognized as key factors in adaptive evolution, there is still much to be learned about how they evolve over time and what genetic mechanisms underlie their important phenotypic effects. In general, our results are consistent with the supergene hypothesis of inversion evolution and suggest that not only are multiple linked loci within the adaptive inversion contributing to its evolution, but that some of those loci may continue to evolve in response to local selective pressures in coastal versus inland habitats. The long-read sequencing assemblies that we report in this study also allowed us to localize three other large chromosomal inversions, each of which illustrates a key component of the evolution of inversions within species. This included the identification of the nested inversion (inv_chr8B) within inv_chr8A, which had not been identified in prior studies. Beyond chromosome 8, we were able to greatly improve the assembly of the inv_chr5A inversion, which was first reported in a cross between inland annual and coastal perennial lines (Holeski et al., 2014). The inv_chr5A inversion is suspected to play a role in adaptation based on elevated allelic differentiation of this region between coastal perennial and inland annual populations (Gould et al., 2018), which is a similar pattern to that for the locally adaptive inv_chr8A inversion. The inv_chr14A inversion was mentioned in Flagel et al. (Flagel et al., 2019) as a 4 MB region of suppressed recombination. We were able to confirm that this region is in fact an inversion. Based on our analyses, the inv_chr14 inversion was less differentiated between the ecotypes, suggesting that it may not be as geographically widespread as the chromosome 5 and 8 inversions. Overall, the results of this study illustrate multiple important aspects of large inversion evolution which we discuss below.

### The evolution of complex nested inversions

The biggest surprise of our study was the discovery of the relatively large inv_chr8B inversion (214-261 kb; depending on genome) nested within the adaptive inv_chr8A inversion. The evolution of nested inversions is not unprecedented, but is also not generally discussed in the context of locally adaptive inversions. Instead, nesting and clustering of multiple inversions in close proximity is typically associated with the evolution of sex chromosomes. For sex chromosomes, the accumulation of successive inversions and other rearrangements leads to contrasting levels of divergence (i.e. strata) for different regions of sex chromosomes, depending on the age of each region of suppressed recombination (D. Charlesworth, 2023; Filatov, 2022; Handley et al., 2004; Olito & Abbott, 2023). This process eventually leads to distinct X/Y or Z/W chromosomes that no longer contain pseudoautosomal regions that recombine at all at meiosis when heterozygous (Bergero & Charlesworth, 2009; D. Charlesworth, 2023). Beyond sex chromosomes, studies in both maize and sunflowers have found evidence for large nested inversions (Dawe, 2022; Mroczek et al., 2006; Todesco et al., 2020). In maize, nested inversions have been important for the evolution of chromosomal knobs that spread by meiotic drive (Dawe, 2022; Mroczek et al., 2006). The significance of the autosomal nested inversions in sunflowers is unclear.

The entire region that includes both inversions on chromosome 8 is known to be involved in local adaptation to coastal perennial versus inland annual habitats (Lowry & Willis, 2010). Given the geographic distribution of the larger inversion (inv_chr8A), it is also likely involved in adaptation to wetter versus drier habitats within inland regions (Lowry & Willis, 2010; Oneal et al., 2014; Twyford & Friedman, 2015). Therefore, natural selection is intimately involved in the spread of inv_chr8A. The question remains as to whether the smaller inv_chr8b inversion also confers habitat-dependent fitness advantages. The inv_chr8B inversion is particularly interesting, as it appears to be an ancient inversion that was later captured by the larger inv_chr8A inversion. These results suggest the intriguing possibility that inv_chr8B is itself an adaptive supergene that now contributes important phenotypic effects to the larger adaptive inv_chr8A supergene.

### The role of context-dependent suppressed recombination in the evolution of adaptive chromosomal inversions

Inversions likely play a major role in adaptation because they suppress recombination in heterokaryotic individuals, while allowing for free recombination in homokaryotic individuals (Berdan et al., 2023; Navarro et al., 1997). This dynamic means that locally adaptive alleles at distant loci on the same chromosome can be maintained in the same haplotypes for populations located in divergent habitats. At the same time, free recombination within each habitat allows for purging of deleterious and maladapted alleles for the same genomic region within habitats (Berdan et al., 2021; B. Charlesworth, 2012; Huang et al., 2022). Further, free recombination in each habitat minimizes the interference of selection to act on multiple loci within that region through Hill-Robertson effects (Felsenstein, 1974; Hill & Robertson, 1966; Roze & Barton, 2006). Our study, as well as a previous one (Gould et al., 2017), found evidence that study’s GA20ox2 has undergone recent selective sweep within the coastal perennial ecotype, while the rest of the inv_chr8A region did not. This suggests that these loci can evolve independently of other parts of the inversion with ecotypes. Likewise, a recent study in sunflowers found that a high level of homozygosity for inversions minimized the mutational load in inverted regions (Huang et al., 2022). In both of these systems, adaptive inversion polymorphisms are thought to primarily occur in a homozygous state in alternative habitats, which facilitates recombination between loci within the inverted region. Overall, the ability of inversions to purge deleterious mutations and minimize Hill-Robertson effects give inversions a major evolutionary advantage over other types of long regions of suppressed recombination, such as pericentromeric regions (Kuhl & Vader, 2019; Wong & Filatov, 2023).

### Candidate genes underlying the adaptive inversion’s phenotypic effects

While the idea that maintenance of adaptive alleles across multiple loci through suppressed recombination is a primary reason for the association of inversions with local adaptation, efforts to identify the genes underlying adaptive phenotypic effects of inversions are still in the initial stages (Berdan et al., 2023). Genes within this inversion involved in the gibberellin (GA) hormone pathway are of particular interest because of the discovery that the addition of gibberellin (GA3) to coastal perennial plants leads them to develop more like inland annual plants by converting vegetative stolons into upcurved flowering branches (Lowry et al., 2019). Introgressing the inland annual orientation of the inversion into the coastal perennial background resulted in nearly identical phenotypic effects on branching and internode elongation as spraying coastal plants with GA3 (Lowry & Willis, 2010). The GA gene, *GA20ox2* is a top candidate gene for contributing to inv_chr8A’s phenotypic effects. Not only were these genes allele frequency outliers in comparisons of perennial and annual populations, but *GA20ox2* was expressed at a higher level in the shoot apex of LMC-L1 than SWB-S1. *GA20ox2* expression is tissue- and time-specific and we were not able to find evidence that it was expressed when analyzing RNA-seq data from leaf tissue (Gould et al., 2017). In other systems (Andrés et al., 2014), *GA20ox2* is typically expressed in the floral shoot apex, similar to our findings in *M. guttatus.* Gould et al (2017) collected tissue only from leaves, which did not detect differential expression of *GA20ox2* in the floral shoot apex. In contrast, we found higher expression of *GA20ox2* in the floral apex for the inland annual than the coastal perennial line when we isolated RNA only from the floral shoot apex tissue. This expression pattern is consistent with the hypothesis that inland annuals flower earlier and have an upright growth architecture due to higher production of gibberellin hormones (Lowry et al., 2019).

While our study, as well as ongoing work, implicates a role for *GA20ox2* in the phenotypic effects of inv_chr8A, the supergene hypothesis requires that we identify multiple genes within the inversion that contribute to adaptive phenotypic divergence. A promising candidate for also contributing to adaptive phenotypic effects is the cluster of R2R3-MYB genes within inv_chr8A. A previous study mapped variation in five vegetative anthocyanin traits to the region inv_chr8A (Lowry et al., 2012), and coastal perennial and inland annual populations differ in anthocyanin pigmentation. While this gene cluster is promising, it has been difficult to link vegetative anthocyanin polymorphisms to evolutionary adaptations (Hatier & Gould, 2008; Hughes et al., 2013; Hughes & Lev-Yadun, 2023; Lee et al., 1979; Manetas, 2006; Schaefer & Rolshausen, 2006). In addition to this R2R3-MYB gene cluster, genes within the inv_chr8B inversion may play a role in the phenotypic effects of the locally adaptive inv_chr8A inversion.

### Little evidence for a role of breakpoint mutation in causing the phenotypic effects of adaptive inversions

While the supergene hypothesis has been championed by evolutionary biologists in recent years, it is not the only mechanism by which inversions can contribute to adaptive phenotypic variation. Inversion mutations can also break genes or alter gene expression through mutations in promoters/enhancers or by changing the local chromatin architecture near breakpoints (Berloco et al., 2014; Elgin & Reuter, 2013; Muller, 1930; Puig et al., 2015; Vogel et al., 2009).

In this study, we found no evidence in this study that the large inversions disrupted any genes. Instead, breakpoints were located primarily in gene-poor, repeat-rich regions of the genome. While we found no evidence that the large inversions in our study broke any genes, it is possible that inversion mutations could have resulted in changing gene expression of nearby genes by disrupting promoter/enhancer elements or changing local chromatin landscapes. Indeed, we did find that many of the genes located in close vicinity of the inversion breakpoints on chromosome 5 and 8 had significantly different levels of transcript abundance between SWB-S1 and LMC-L1 plants. From our study, we cannot establish whether the breakpoint mutation itself caused these differences in gene expression or if that regulatory divergence evolved later. A recent study in *Drosophila* that created synthetic inversions at the position of naturally occuring inversion polymorphism did not cause any major effects on gene expression. (Kapun et al., 2023; Said et al., 2018) found that no greater elevation in gene expression for genes near the breakpoints of the Drosophila In(3R)Payne inversion versus elsewhere in the inversion. These results collectively suggest that the evolution gene expression divergence for inversions often happens over time after the inversion mutation through the gradual accumulation of new mutations.

## Authors contribution

L.M.K, D.B.L, and C.E.N designed research; L.M.K., C.E.N, L.E.S. and S.K.K.R performed research, and L.M.K. and C.E.N. analyzed data; and L.M.K., C.E.N., L.E.S., and D.B.L. wrote the paper.

## Data, materials, and software availability

Code for genome assembly can be found at https://github.com/niederhuth/mimulus-assembly and code for calling structural variants and SNPs, reanalysis of pooled sequencing and RNAseq, and figures can be found at https://github.com/Kollarlm/Breakpoint-mutations-supergene-effects-and-ancient-nested-rearrangements-in-yellow-monkey-flower.git. Raw ONT sequencing files are available through the NCBI Sequence Read Archive under project identifier PRJNA1047892 – accessions SAMN38597945 (LMC-L1) and SAMN38597946 (SWB-S1). Raw RNA sequencing files are also available through the NCBI Sequence Read Archive under project identifier PRJNA1054563. The genome assemblies, annotations, TE annotations, and other supporting documents can be found on Dryad under DOI:10.5061/dryad.2ngf1vhwj and CoGE XXX.

## Acknowledgments

This study was funded by the National Science Foundation (NSF) Plant Genome Research Program 2010769 to LMK and 2109560 to LES, Michigan State University Plant Biology start-up funds to CEN, and by two National Science Foundation Division of Integrative Organismal Systems Grants to DBL (IOS-1855927 and IOS-2153100). We would like to thank Oxford Nanopore Technologies and the MSU Genomics Core for providing reagents and services that generated sequencing data that made this work possible. We would also like to thank Kevin Childs for guidance on genome annotation, Eleanore Ritter for guidance in detecting structural variants, and Lane Vitek for lab work.

