## supplemental for "The evolutionary dynamics of locally adaptive chromosome inversions in *Mimulus guttatus*"

***SI Appendix***

| Metrics | SWB-S1 | LMC-L1 | IM62.v2 |
| --- | --- | --- | --- |
| Nanopore Reads | 9,423,361 reads | 9,255,001 reads | - |
| Base Pairs | 101.76 Gbps | 93.35 Gbps | - |
| Assembly Size | ~278 Mbps | 277 Mbps | ~290 Mbps |
| Contig Number | 707 | 1198 | - |
| N50/N90 Contigs | ~4.90/~0.72 Mbps | ~5.83/~0.49 Mbps | - |
| Sequence in Chromosomes | 93.70% | 93.00% | 93.40% |
| N50/N90 Scaffolded | ~18.8/~12.0 Mbps | ~18.2/~11.7 Mbps | ~21.2/13.6 Mbps |
| Genes | 26,876 | 27,583 | 28.14 |
| BUSCO | 98.00% | 98.20% | 96.90% |
| LAI | 13.47 | 16.28 | 8.79 |

**Table S1.** Genome assembly statistics for LMC-L1 and SWB-S1 genomes with comparisons of IM62 v2 where applicable.

| Location | LMC-L1  specific  genes | SWB-S1  specific  genes | Shared  genes |
| --- | --- | --- | --- |
| Genome | 5,347 | 4,797 | 19,245 |
| inv_chr5A | 113 | 90 | 369 |
| inv_chr8B | 9 | 8 | 36 |
| inv_chr8A (including small inversion) | 177 | 125 | 613 |
| inv_chr8A (large inversion without small inversion) | 168 | 117 | 577 |
| inv_chr14A | 247 | 308 | 1,133 |

**Table S2.** The number of shared and ecotype- specific genes for the 4 large chromosomal inversions and genomewide. These genes were identified through orthology constrained synteny.

| Ecotype | Inversions | Deletion | Duplications | Insertion | SVs within genes |
| --- | --- | --- | --- | --- | --- |
| LMC-1 | 266 | 3,752 | 214 | 4,174 | 12,013 |
| SWB-S1 | 253 | 4,022 | 134 | 3,949 | 11,582 |

**Table S3.** The number of structural variants between LMC-L1 and SWB-S1 and the number of SVs that fall within genic regions.

| Location | Variant type | SWB-S1 total SNPs | SWB-S1 number of genes affected | LMC-L1 total SNPs | LMC-L1 number of genes affected |
| --- | --- | --- | --- | --- | --- |
| Genome - wide | Stop gain | 2,787 | 2,337 | 2,638 | 2,234 |
| inv_chr5A | Stop gain | 71 | 58 | 61 | 55 |
| inv_chr8B | Stop gain | 10 | 8 | 8 | 5 |
| inv_chr8A | Stop gain | 99 | 82 | 94 | 78 |
| inv_chr14A | Stop gain | 32 | 25 | 25 | 21 |
| Genome - wide | Stop loss | 723 | 696 | 700 | 676 |
| inv_chr5A | Stop loss | 19 | 18 | 14 | 14 |
| inv_chr8B | Stop loss | 4 | 4 | 2 | 2 |
| inv_chr8A | Stop loss | 34 | 32 | 32 | 29 |
| inv_chr14A | Stop loss | 11 | 10 | 8 | 8 |
| Genome - wide | Frameshift deletion | 7,299 | 4,592 | 7,541 | 4,669 |
| inv_chr5A | Frameshift deletion | 182 | 117 | 177 | 115 |
| inv_chr8B | Frameshift deletion | 17 | 10 | 14 | 11 |
| inv_chr8A | Frameshift deletion | 291 | 181 | 282 | 171 |
| inv_chr14A | Frameshift deletion | 66 | 45 | 77 | 50 |
| Genome - wide | Frameshift insertion | 7,002 | 4,449 | 6,951 | 4,431 |
| inv_chr5A | Frameshift insertion | 176 | 118 | 174 | 112 |
| inv_chr8B | Frameshift insertion | 15 | 12 | 21 | 10 |
| inv_chr8A | Frameshift insertion | 283 | 170 | 277 | 169 |
| inv_chr14A | Frameshift insertion | 76 | 47 | 56 | 44 |
| Genome - wide | Splicing | 3,986 | 3,311 | 3,392 | 4,109 |
| inv_chr5A | Splicing | 93 | 81 | 83 | 117 |
| inv_chr8B | Splicing | 12 | 9 | 13 | 18 |
| inv_chr8A | Splicing | 165 | 130 | 129 | 152 |
| inv_chr14A | Splicing | 35 | 29 | 42 | 52 |

**Table S4.** LMC-L1 and SWB-S1 SNPs called from WGS were annotated and filtered to include stop gain, stop loss, frameshift deletion, frameshift insertion, and splicing variants.

| Location | LMC-L1  size | SWB-S1  size | LMC-L1  SNPs | SWB-S1  SNPs |
| --- | --- | --- | --- | --- |
| Genome |  |  | 6,192,716 | 6,234,953 |
| inv_chr5A | 4,196,511 bp | 4,010,835 bp | 141,267 | 128,725 |
| inv_chr8B | 213,792 bp | 261,302 bp | 8,606 | 7,732 |
| inv_chr8A  (including small inversion) | 6,754,390 bp | 5,606,574 bp | 223,542 | 181,793 |
| inv_chr8A  (without small inversion) | 6,745,784 bp | 5,598,842 bp | 214,936 | 174,061 |
| inv_chr14A | 2,415,580 bp | 2,784,925 bp | 63,711 | 79,338 |

**Table S5.** The number of filtered SNPs within large chromosome inversions. Note, while we are unsure if we have captured the entire chromosome 5 inversion due to its proximity near the end of the chromosome, we take caution when interpreting differences for this particular inversion.

| Inversion | | LMC-L1 Start | SWB-S1  Start | LMC-L1  End | SWB_S1  End | LMC-L1  Genes | SWC-S1  Genes | LMC-L1  Pseudogenes | SWB-S1  Pseudogenes |
| --- | --- | --- | --- | --- | --- | --- | --- | --- | --- |
| inv_chr5A | 13,650,670 | | 14,125,309 | 17,847,181 | 18,136,144 | 535 | 501 | 54 | 43 |
| inv_chr8A | 850,429 | | 858,736 | 7,604,769 | 6,465,310 | 878 | 819 | 90 | 74 |
| inv_chr8B | 1,032,334 | | 5,998,986 | 1,246,126 | 6,260,288 | 50 | 45 | 6 | 8 |
| inv_chr14A | 5,329,939 | | 5,892,556 | 7,791,197 | 8,677,481 | 281 | 294 | 22 | 19 |

**Table S6.** Chromosome 5, 8, and 14 inversion breakpoints according to synteny and number of genes. There were a total of 27,583 detected in LMC-L1 and 26,876 genes in SWB-S1. Note chromosome 8 values include the smaller inversion located within it.

| Gene | Genome | Function | Differentially.expressed |
| --- | --- | --- | --- |
| MgL1_05g16120 | LMC-L1 | HXXXD-type acyl-transferase family proteins | Between genotypes and  between field sites within each ecotype |
| MgL1_08g01750 | LMC-L1 | Major facilitator superfamily protein | Between genotypes, G x E, and between field sites within coastal perennials |
| MgL1_14g10910 | LMC-L1 | Triosephosphate isomerase | Between genotypes and  between field sites within each ecotype |
| MgS1_05g10260 | SWB-S1 | HXXXD-type acyl-transferase family proteins | G x E |
| MgS1_08g08930 | SWB-S1 | Unknown function | Between field sites within inland annuals and G x E |
| MgS1_14g08050 | SWB-S1 | Unknown function | Between genotypes and between field sites within each ecotype |
| MgS1_14g08070 | SWB-S1 | Ctr copper transporter family | Between genotypes and between field sites within each ecotype |

**Table S7.** Differentially expressed genes at the inversion breakpoints.

**Figure supplemental figures**

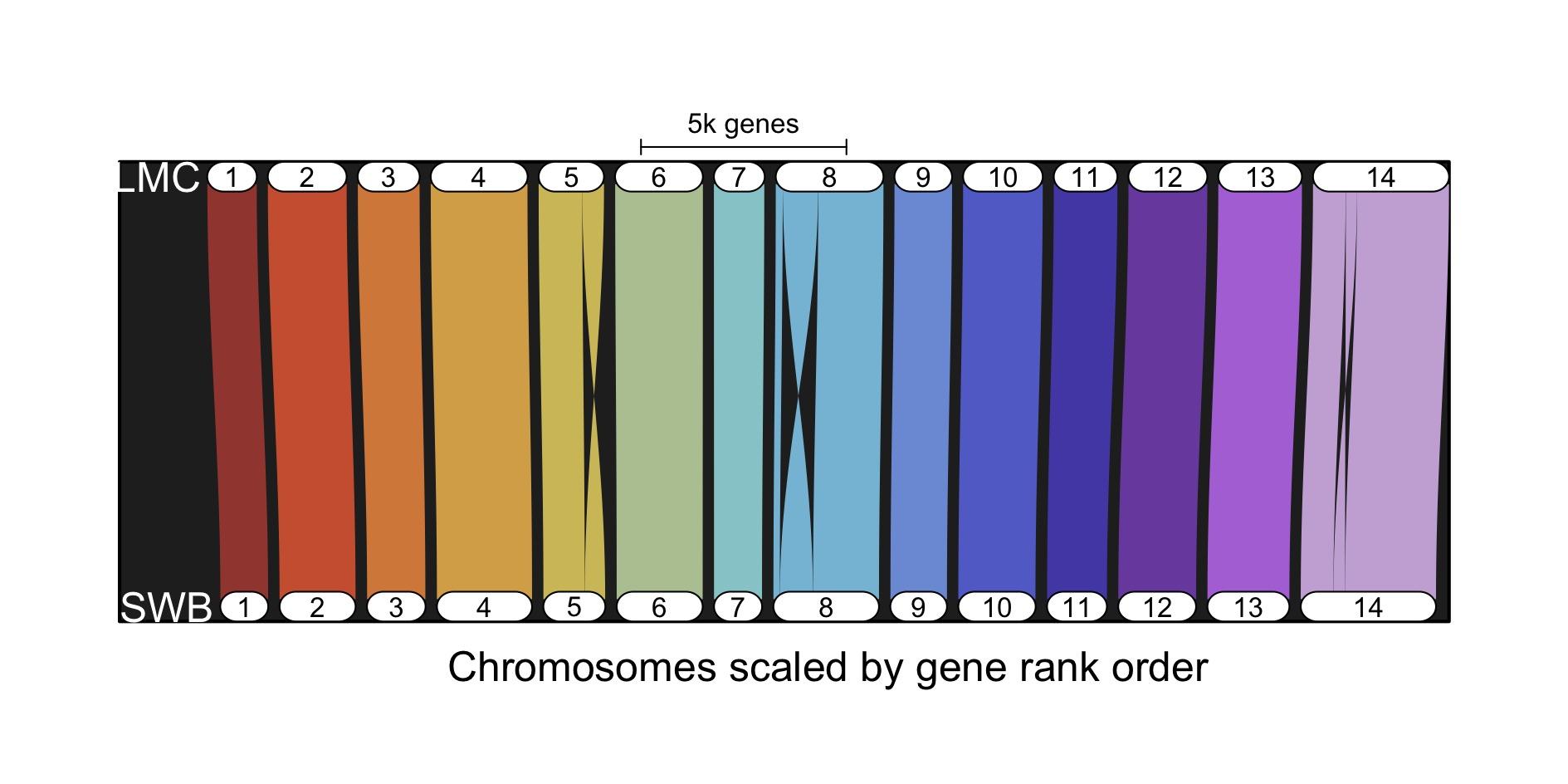

**Fig. S1.** Comparison of LMC-L1 and SWB-S1 using synteny constrained orthology (GENESPACE). Chromosomes 5, 8, and 14 show the chromosome inversions.

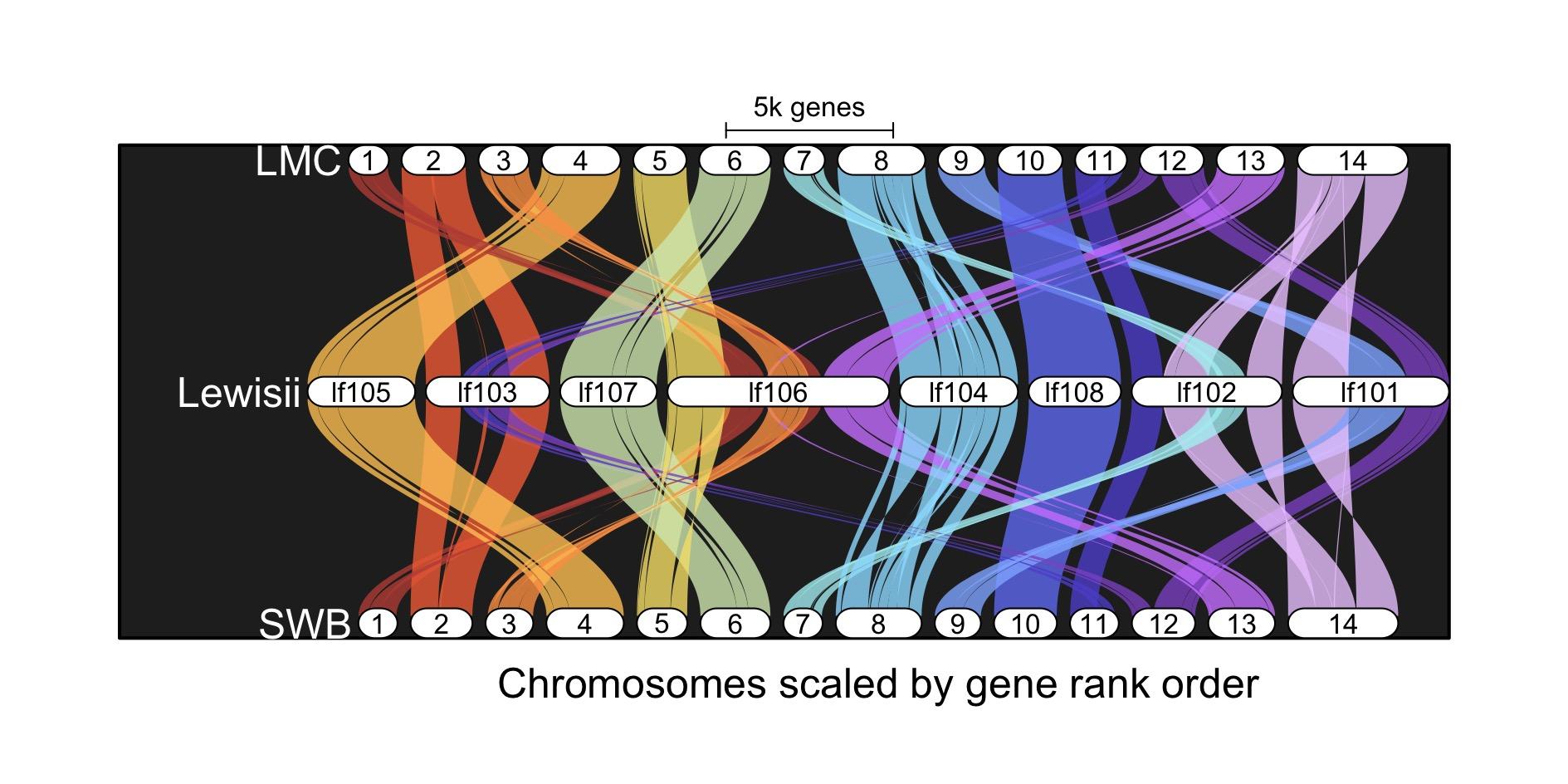

**Fig. S2.** Comparison of LMC-L1, SWB-S1, and Lewisi genomes using synteny constrained orthology (GENESPACE). Chromosomes 5, 8, and 14 show *Lewisi* and LMC-L1 share the same orientation for those inversions. This suggests these inversions arose in LMC-L1 first.

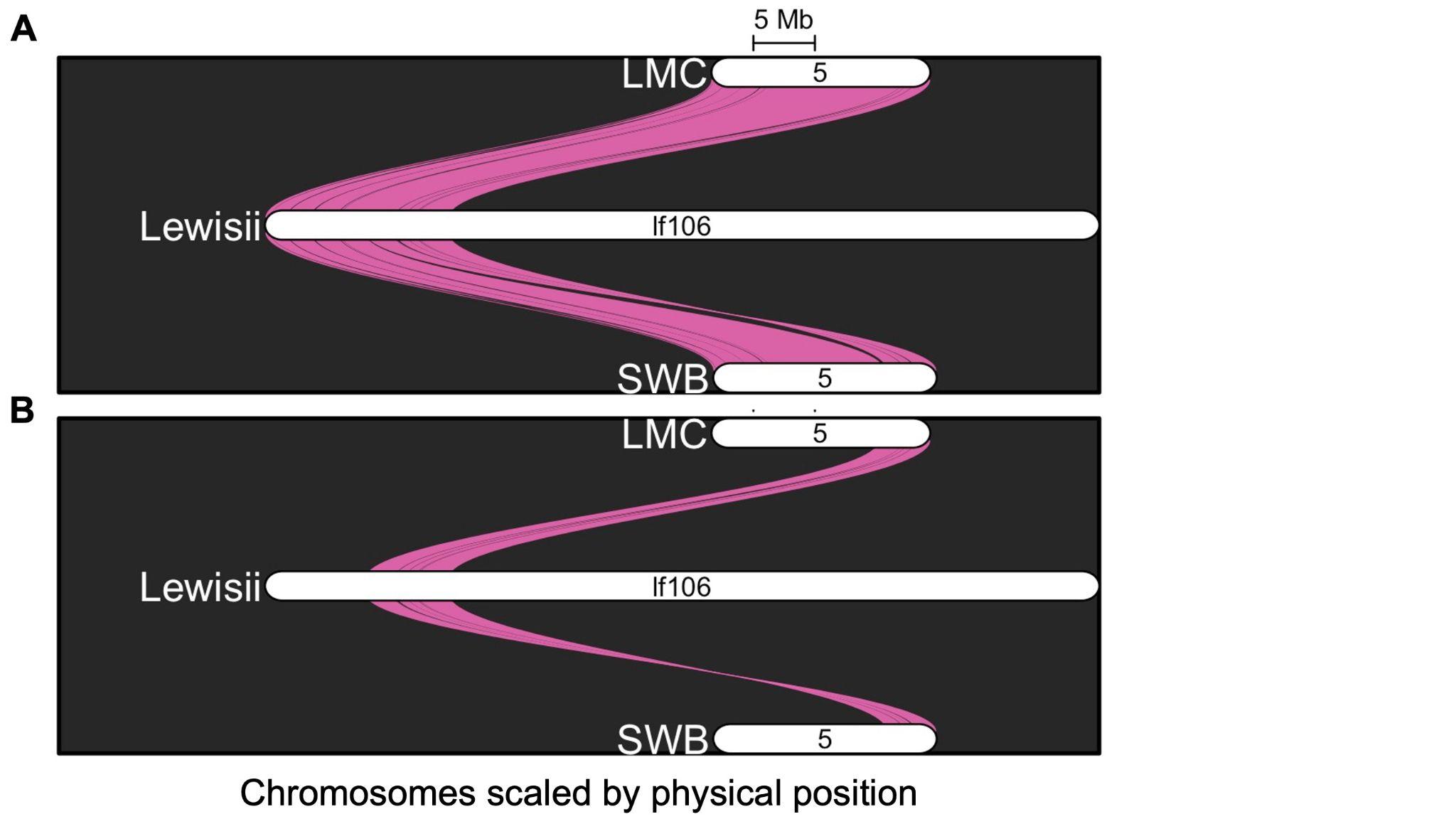

**Fig. S3.**Comparison of LMC-L1, SWB-S1, and Lewisi inv_chr5A using synteny constrained orthology (GENESPACE). **A** represents the entire chromosome 5 while **B** represents inv_chr5A. Inv_chr5A in *Lewisi* and SWB-S1 share the same orientation suggesting these inversions arose in LMC-L1 individuals.

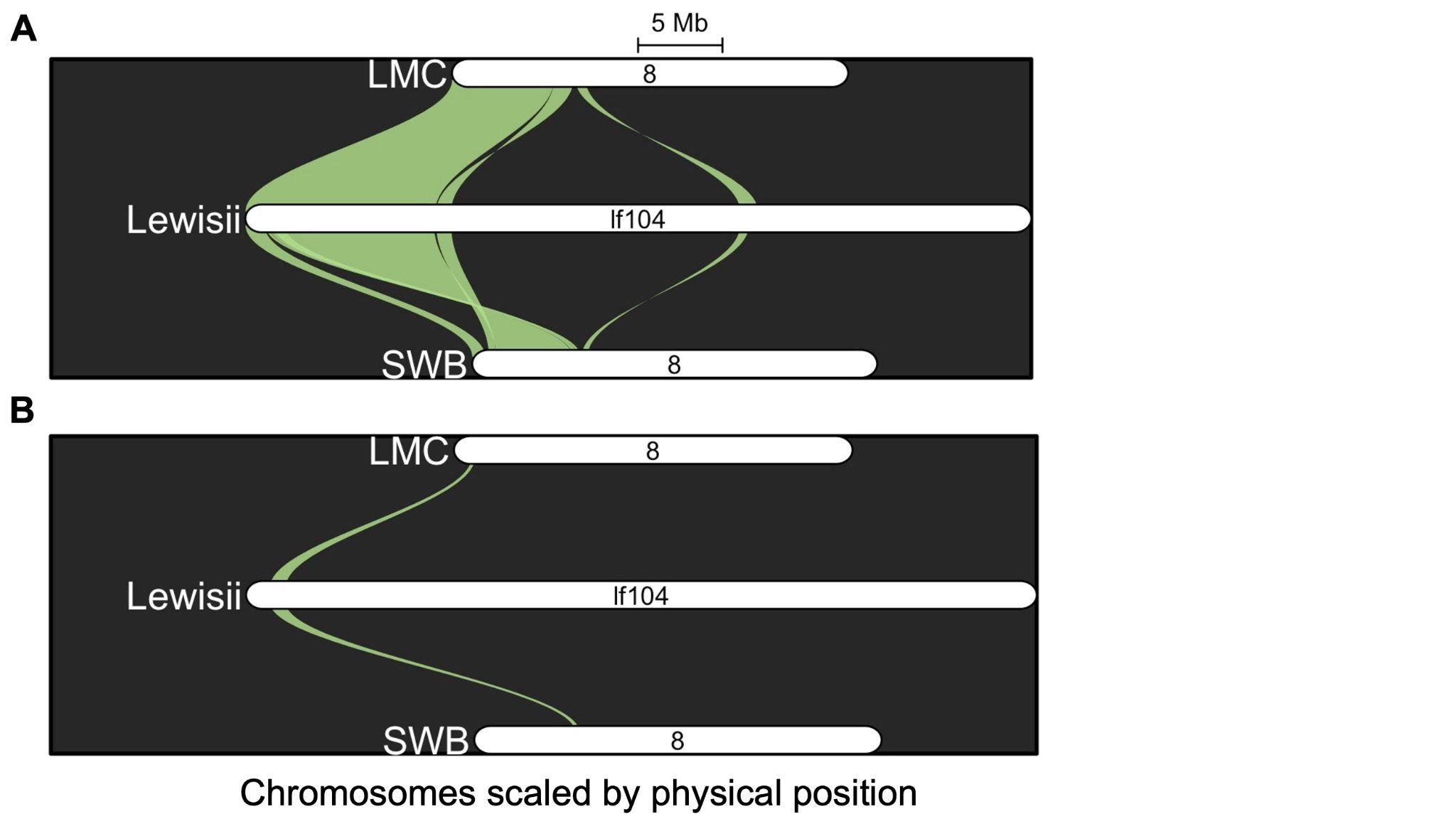

**Fig. S4.** Comparison of LMC-L1, SWB-S1, and Lewisi internal chromosome 8 inversion using synteny constrained orthology (GENESPACE). **A** represents the inv_chr8A while **B** represents inv_chr8B. Inv_chr8B in *Lewisi,* LMC-L1, and SWB-S1 share the same orientation suggesting these inversions arose in SWB-S1 individuals.

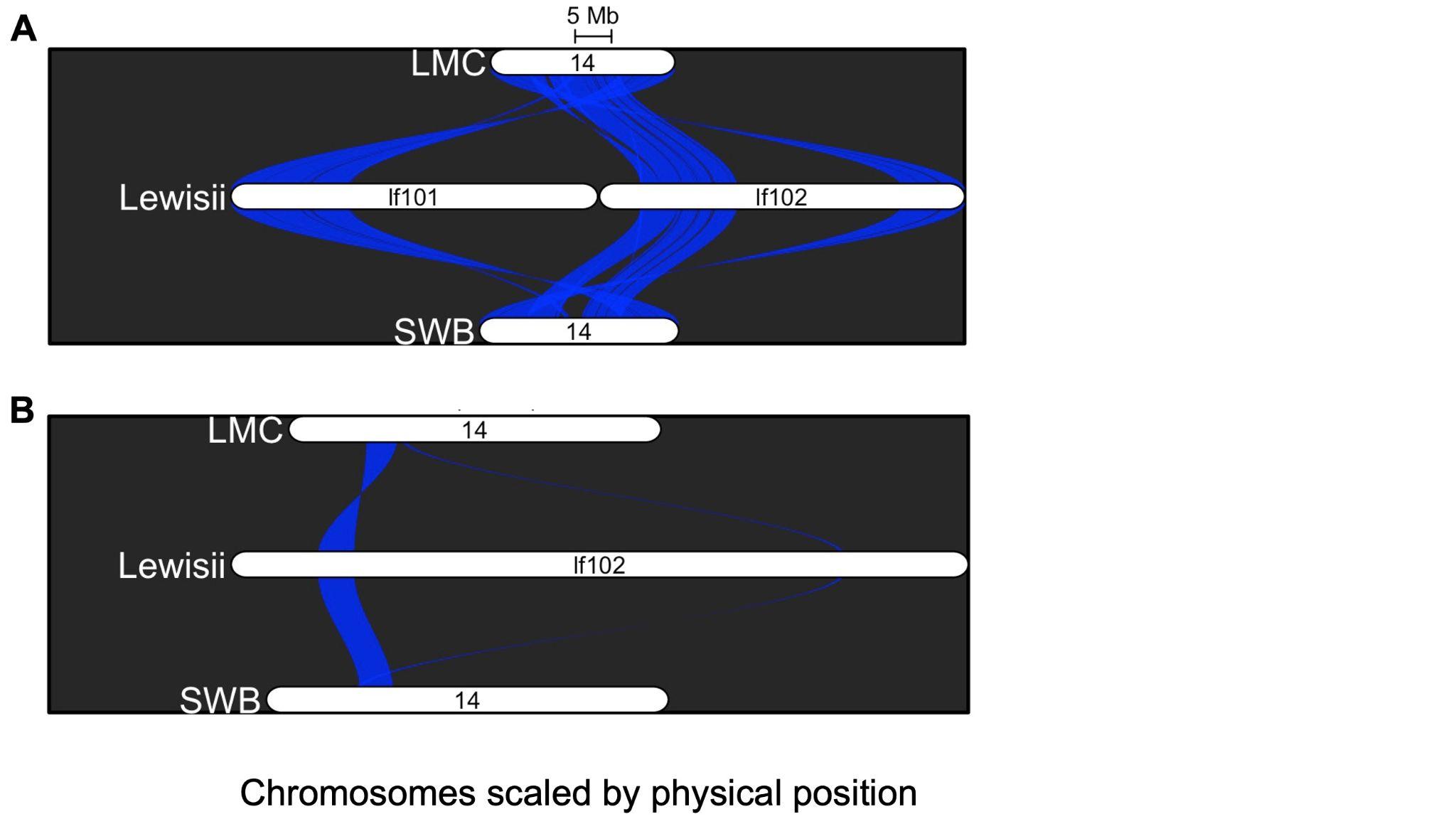

**Fig. S5.** Comparison of LMC-L1, SWB-S1, and Lewisi inv_chr14A using synteny constrained orthology (GENESPACE). **A** represents the entire chromosome 14 while **B** represents inv_chr14A. Inv_chr14A in *Lewisi* and SWB-S1 share the same orientation suggesting these inversions arose in LMC-L1 individuals.

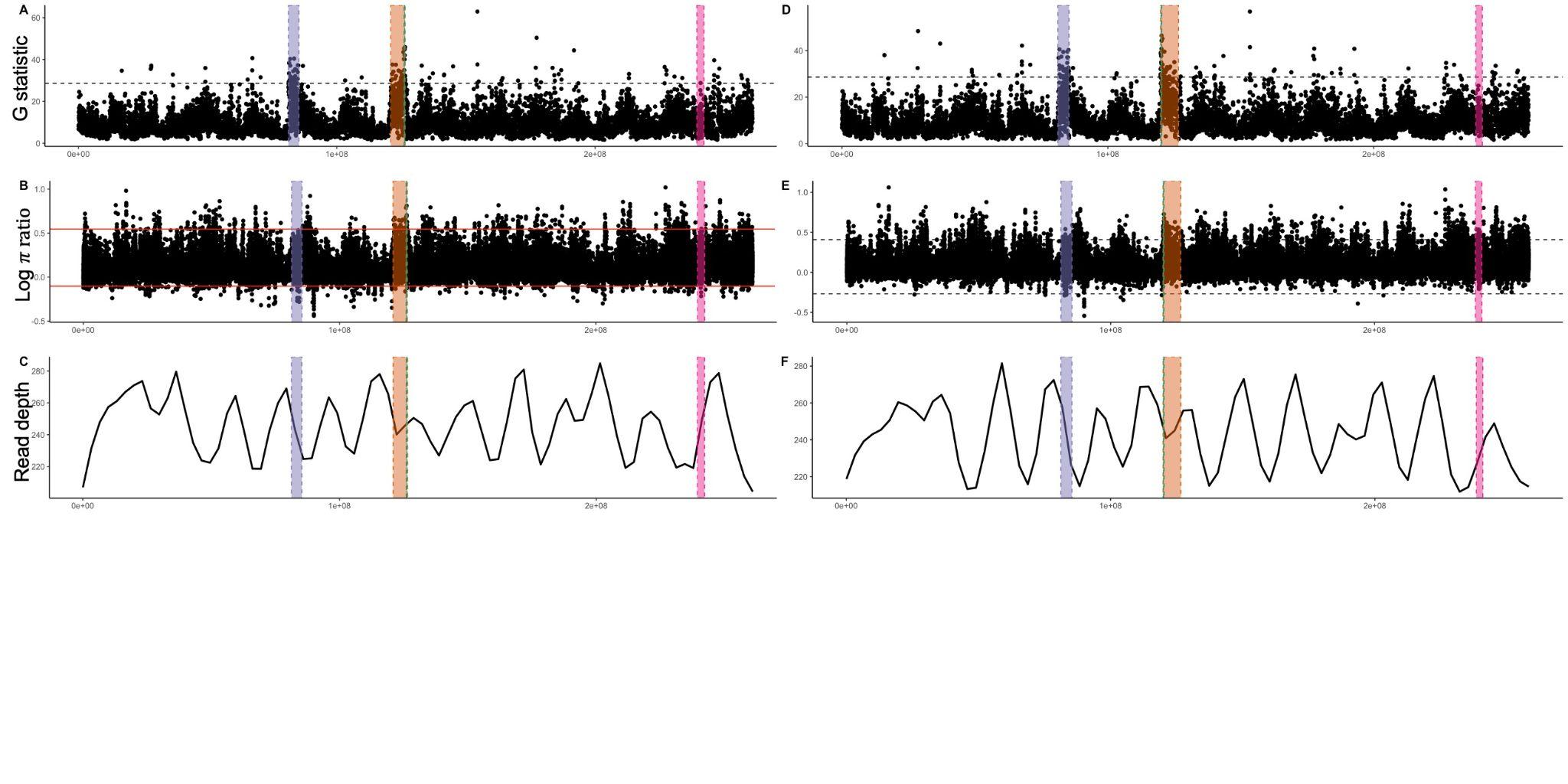

**Fig. S6.** Estimates of *G* (A - SWB-S1 and D - LMC-L1), π ratio (B - SWB-S1 and E - LMC-L1) , and read depth (C - SWB-S1 and F - LMC-L1) across the SWB-S1 (left) and LMC-L1 (right) genome when both coastal and inland pools were aligned. The purple region represents the chromosome 5 inversion, the orange region represents the chromosome 8 inversion with the smaller inversion in green, and the chromosome 14 inversion is represented in π nk.

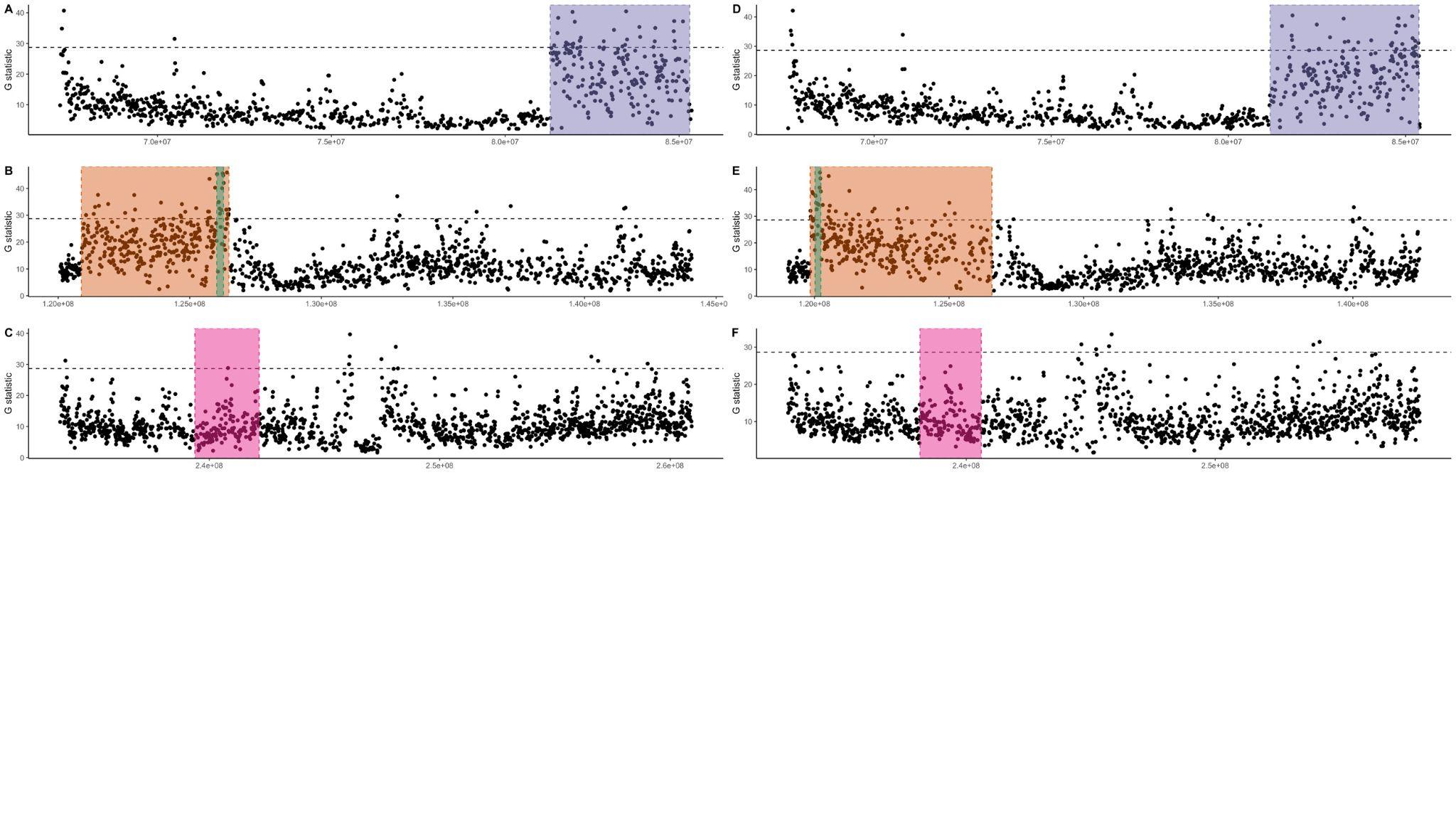

**Fig. S7.** Estimates of *G* statistic when aligned to the SWB-S1 (right A - C) and LMC-L1 (right D - E) genomes. **A** and **D** represent chromosome 5 with the purple region highlighting the inversion. **B** and **E**  represent chromosome 8 with the large inversion highlighted in orange and the smaller inversion highlighted in green. **C** and **F** represent chromosome 14 with the inversion highlighted in π nk.

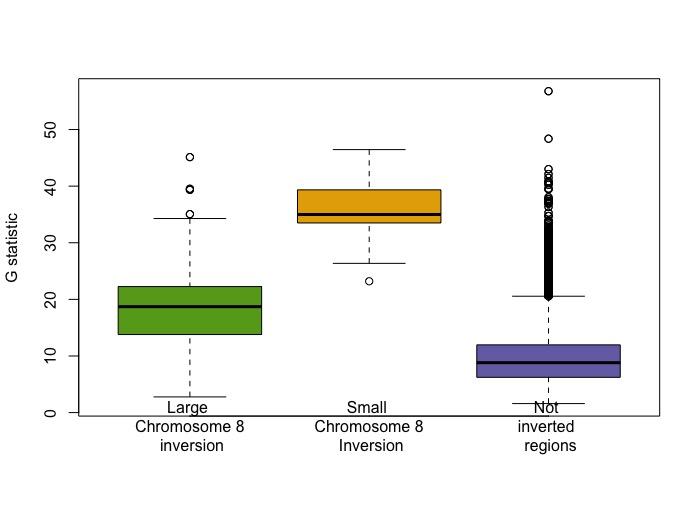

**Fig. S8.** *G* statistic windows using the LMC-L1 genome were separated into the large chromosome 8 inversion excluding windows falling within the small chromosome 8 inversion, smaller chromosome 8 inversion, and non-inverted regions along chromosome 8. We plotted the ANOVA (G statistic ~ region of window). The results from the ANOVA showed a significant difference in *G* for the different regions along chromosome 8 (d.f. = 2, p < 2e-16)

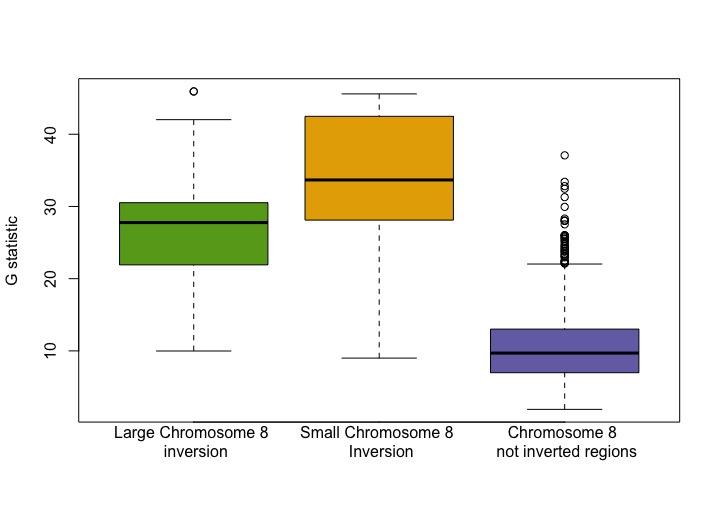

**Fig. S9**. *G* statistic windows using the SWB-S1 genome were separated into the large chromosome 8 inversion excluding windows falling within the small chromosome 8 inversion, smaller chromosome 8 inversion, and non-inverted regions along chromosome 8. We plotted the ANOVA (*G* statistic ~ region of window). The results from the ANOVA showed a significant difference in *G* for the different regions along chromosome 8 (d.f. = 2, p < 2e-16)

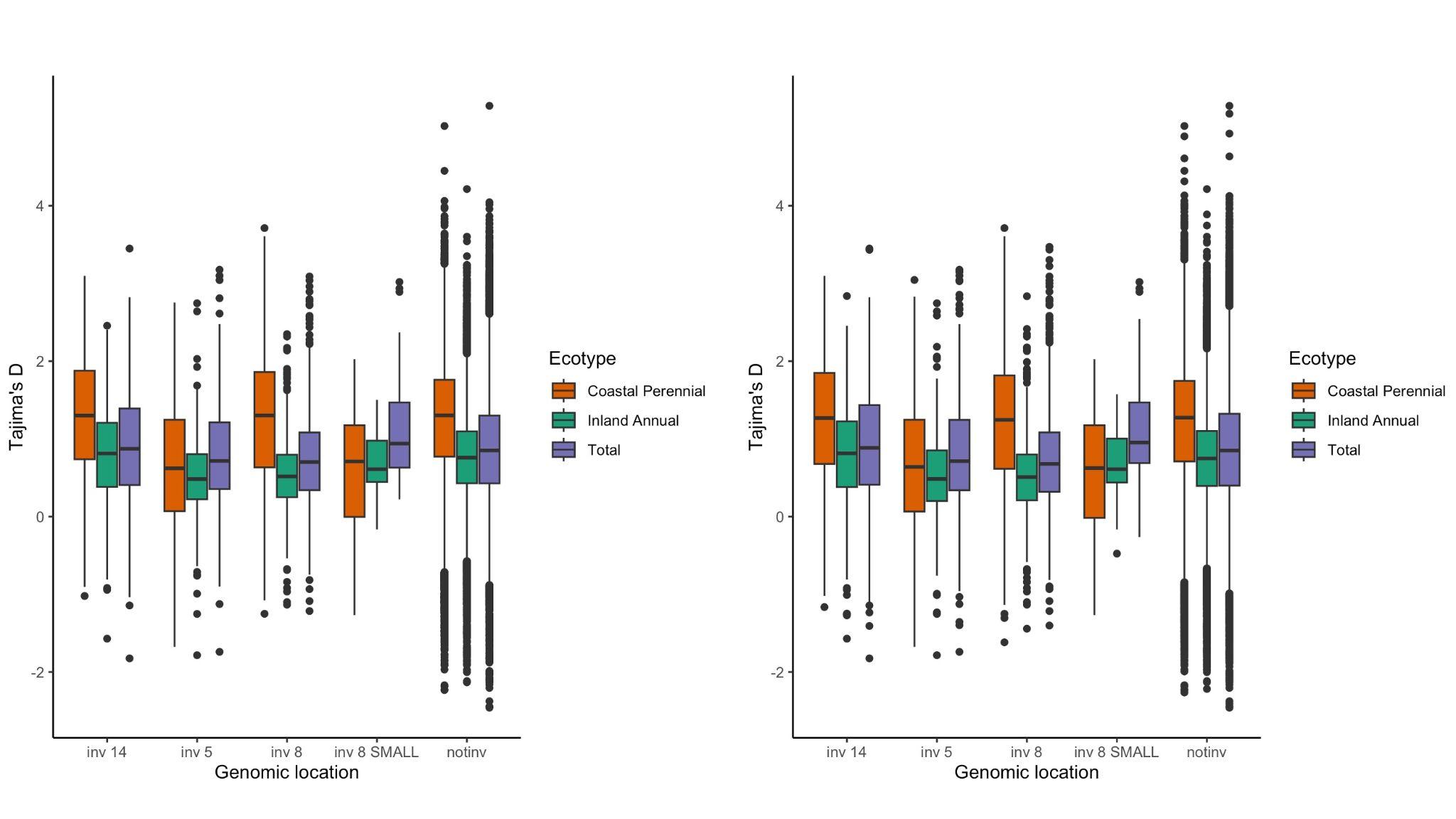

**Fig. S10.** Estimates of Tajima’s D when aligned to the SWB-S1 genome. In **A** we included both syntenic and nonsyntenic genes whereas in **B** we filtered out nonsyntenic genes to see if there were differences driven by nonsyntenic genes.

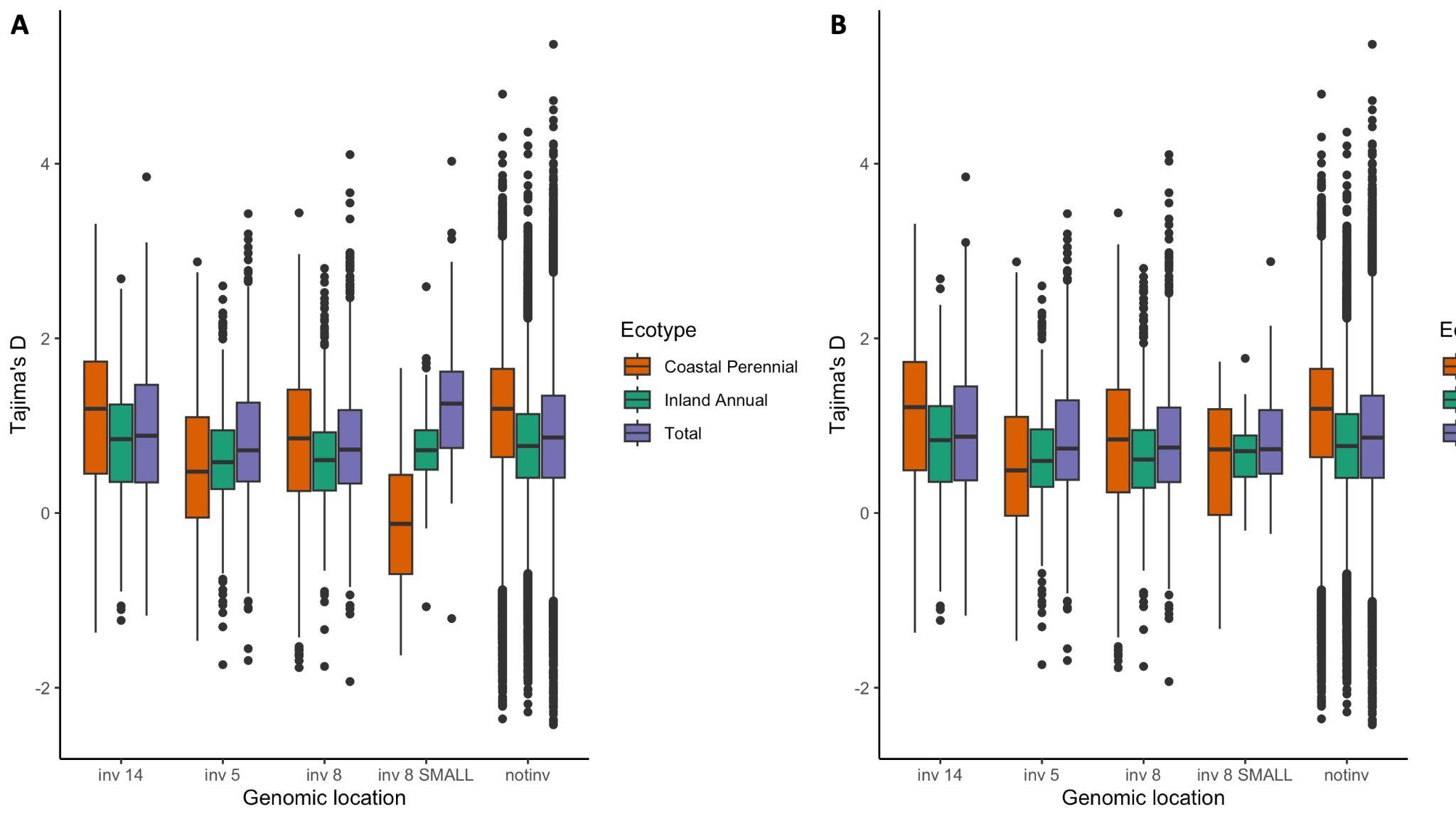

**Fig. S11.** Estimates of Tajima’s D when aligned to the LMC-L1 genome. In **A** we included both syntenic and nonsyntenic genes whereas in **B** we filtered out nonsyntenic genes to see if the negative Tajima’s *D*  was caused by nonsyntenic genes. It appears that there are differences in gene content in the small chromosome 8 inversion that drives these differences in *D*.

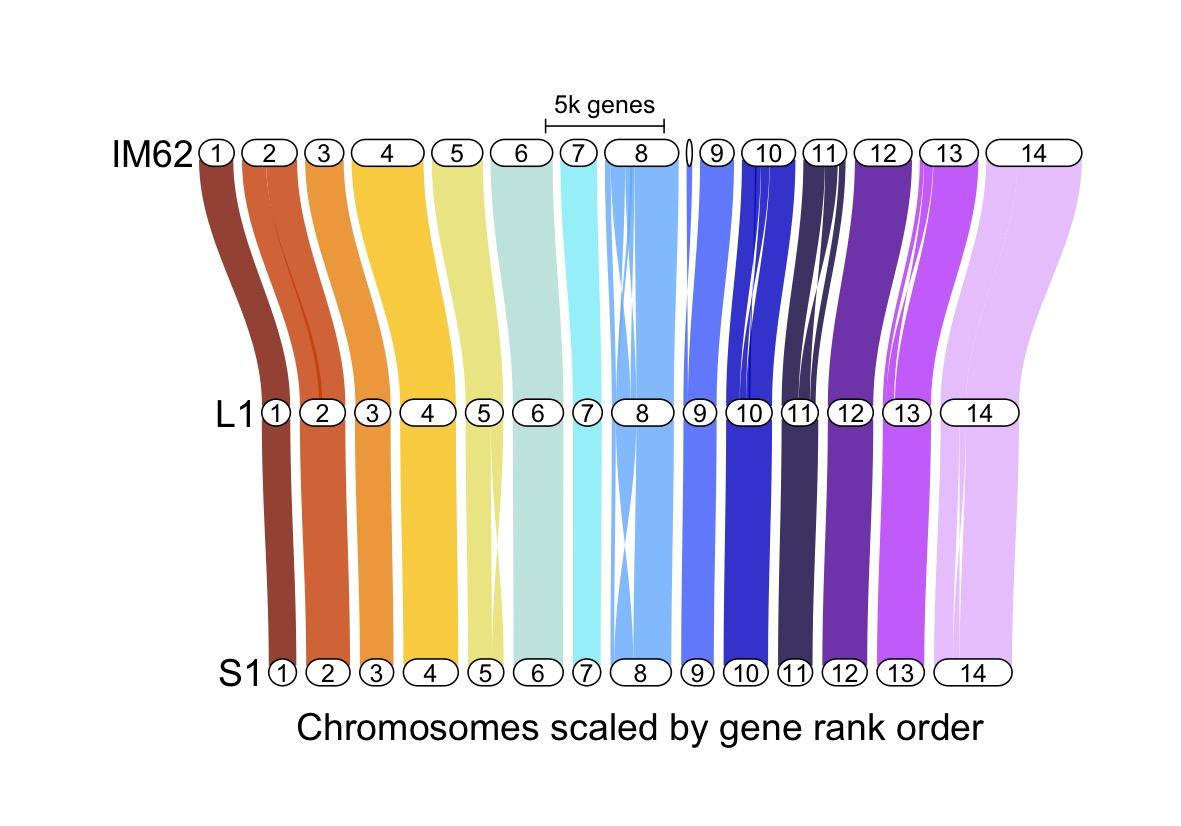

**Fig. S12.** GENESPACE (orthology constraint synteny) plot comparing the LMC-L1, SWB-S1, and IM62.

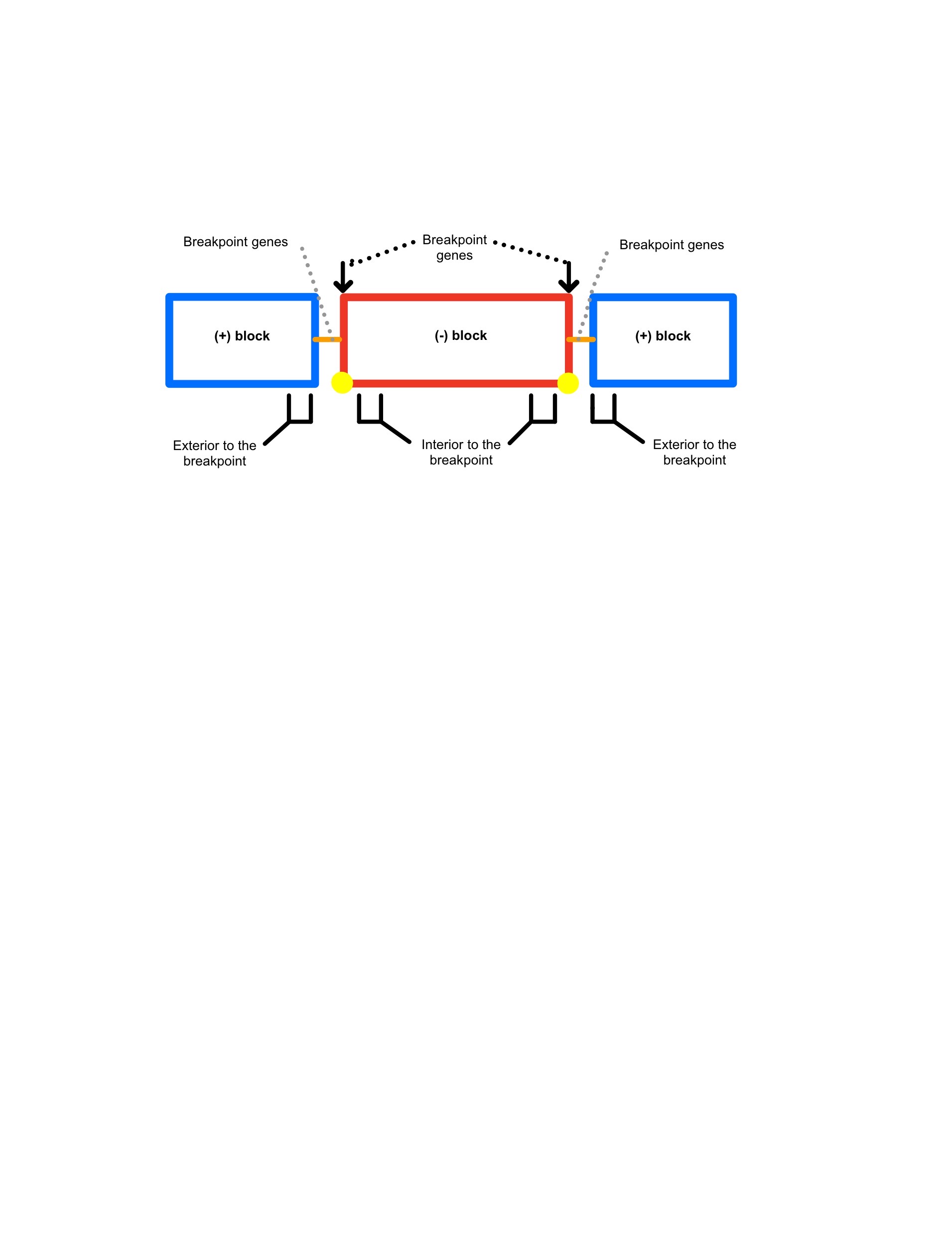

**Figure S13.** The blue blocks represent non-inverted synthetic blocks whereas the red block represents the inversion represented by inverted (-) block(s). The yellow circles signify the first and last gene of the syntenic block and thus the inversion breakpoints. Occasionally, genes that did not fall in a syntenic block were found between the inverted and non inverted blocks. Those genes, represented by the orange lines, were also considered breakpoint genes. Genes interior to the breakpoints are the first genes upstream or downstream of the first or last gene of the inverted block. Genes exterior to the breakpoints were the last gene of the non inverted syntenic block upstream of the inverted block and the first gene of the non inverted syntenic gene downstream of the inverted block.

**SI Appendix, Supplementary methods**

**Supplementary methods 1: Read based alignment structural variant calling methods**

We aligned the quality trimmed ONT reads to the pseudo chromosome scale reference genome using minimap2 (v2.24;(Li, 2018)). Following alignment, we called structural variants (SVs) using sniffles (Sedlazeck et al., 2018), svim (Heller & Vingron, 2019), and pbsv. We filtered the resulting vcf files to include only passing homozygous SVs greater than 50 bps in length. The final output only included inversions, duplications, insertions, and deletions. We merged the vcf files from the three structural variant callers with the program survivor (Jeffares et al., 2017).

**Supplementary methods 2: Gould et al 2017, estimating SNP depleted regions of the genome methods**

To determine regions of depleted SNPs, we incorporated sequencing of coastal perennial (101 accessions) and inland annual (92 accessions) pools from Gould et al (Gould et al., 2017). Each pool was sequenced across two lanes of v2 Rapid lanes on the Illumina High-Seq 2500 platform. To analyze this data with our new genomes, we followed the bioinformatics pipeline published in Gould et al (Gould et al., 2017). We used Trimmomatic (Bolger et al., 2014) to quality-trim forward and reverse reads to a minimum length of 50 bp with a phred score threshold of 33 and removed unpaired reads using SAMTOOLs v1.19.2 (Danecek et al., 2021). Filtered reads were aligned using BWA-mem2 v2.2.1 (Vasimuddin et al., 2019) to our inland annual (LMC) and coastal perennial (SWB) genomes, retaining only uniquely mapped reads. We removed PCR duplicates using PICARDTOOLS’ *MarkDuplicates* function. We generated a pileup file using SAMTOOLS v1.19.2 (Danecek et al., 2021) and called SNPs using SNAPE-pooled (Raineri et al., 2012). We used an informative prior on the allele frequency spectrum and priors for theta and divergence were both set to 0.05 as noted in (Gould et al., 2017). In order to compare our results to (Gould et al., 2017) we closely followed their workflow and used their cited scripts (<https://bitbucket.org/billiegould/genomics_tools/src/master/SNAPEtools/>). We filtered bases based on the probability of a SNP >= 95% (variant) or <= 5% SNP (invariant reference site) while all other sites in between those cutoffs were considered missing data. We removed bases with less than 50x coverage in either pool. For the following statistics, we split the genome into 1000 bp windows and removed windows with an average depth of coverage greater than 2 standard deviations away from the mean (mean = 246.52, standard deviation = 128.03). Using scripts from (Gould et al., 2017), we calculated windowed statistics for Fst, G, and the ratio of inland annual (IA) to coastal perennial (CP) nucleotide diversity ( π IA/ π CP). We focused on the top 1% of G and FST and the top and bottom 1% of π IA/ π CP. We used BEDtools *intersect* (Quinlan & Hall, 2010) to identify genes and promoters (1000 bp upstream of the transcription start site) containing windows flagged for FST, G, and π IA/ipCP outliers. Lastly, we calculated Tajima’s *D* statistic for genes with less than 50% missing data.

**Supplementary methods 3: RNA-seq Analysis**

To evaluate patterns of transcript expression for inversion genes, we reanalyzed RNA-seq data sets from Gould et al. (Gould et al., 2018) in the context of our newly assembled genomes. These data included RNA samples from a field reciprocal transplant experiment, where LMC-L1 and SWB-S1 individuals were grown at both inland and coastal field sites (Gould et al., 2018). The tissue collected only includes leaf tissue and thus, our analysis of differentially expressed genes is limited to leaf tissue. Sequence data were mapped with STAR (Dobin et al., 2013) using a 2-pass mapping method to both the LMC-L1 and SWB-S1 reference genomes.

We used DESeq2 (Love et al., 2014) to determine differential gene expression between genotypes and between genotypes at differing field sites. We included genotype, field site (inland or coastal), and the interaction of genotype and field site as fixed effects for each gene. We calculated a Wald test statistic per gene to find significant differences. P-values were corrected using DESeq2’s default settings, which incorporates Benjamini- Hochberg corrections and FDR <= 0.1 for a gene to be identified as differentially expressed. Additionally, we applied a log2 fold change cutoff of 0.05.
